# Circadian photoreceptor CRYPTOCHROME promotes wakefulness under short winter-like days via a GABAergic circuitry

**DOI:** 10.1101/2023.10.02.560507

**Authors:** Lixia Chen, Danya Tian, Chang Su, Luoying Zhang

**Author notes:** Correspondence: Luoying Zhang.

## Abstract

A cardinal symptom of seasonal affective disorder (SAD, also known as winter depression) is hypersomnolence, while the cause of this “winter sleepiness” is not known. Here we found that lack of the circadian photoreceptor *cryptochrome* (*cry*) leads to increased sleep under short winter-like days in fruit flies, reminiscent of the hypersomnolence in SAD. CRY functions in neurons that synthesize the major inhibitory neurotransmitter GABA, including the small ventral lateral neurons which are known to be circadian pacemakers, and down-regulates the GABAergic tone. This in turn leads to increased neural activity of the wake-promoting large ventral lateral neurons, a subset of circadian neurons that are inhibited by GABA-A receptor. CRY protein is known to be degraded by light, thus rendering CRY to be functional within this GABAergic circuitry to enhance wakefulness only under short day length. Taken together, we demonstrate a mechanism that specifically regulates wakefulness under short winter-like days, which may provide insights regarding the winter sleepiness in SAD.

## Introduction

Seasonal affective disorder (SAD), also known as winter depression, is characterized by the onset of depression in fall/winter months and spontaneous remission in the spring/summer (Rosenthal et al., 1984). It is generally agreed that SAD is caused by lack of day light in the fall/winter months due to the short day length (or photoperiod), as bright light therapy is effective and most commonly used for treating the depression associated with SAD (Golden et al., 2005). The prevalence of SAD is ∼1-10% world-wide, with symptoms lasting for approximately 40% of the year (Kurlansik and Ibay, 2012). About 64-80% of SAD patients report a winter increase in sleep, ranging from 30 min to 2 h longer in duration compared to controls, which is considered to be a distinguishing symptom in the characterization and diagnosis of SAD (Wescott et al., 2020). Currently almost nothing is known regarding the underlying mechanism of this winter hypersomnolence.

This phenomenon of winter hypersomnolence in SAD patients implicates distinct mechanisms that regulate sleep under short winter-like photoperiods vs. longer non-winter-like photoperiods, as the sleep of these individuals appear to be selectively perturbed under short photoperiods. However, the mechanisms by which sleep duration is determined under different photoperiods have not been characterized. Since the circadian clock is believed to be important for adaptations to seasonal changes in the environment and in particular, seasonal changes of photoperiod, we hypothesize that the circadian clock may also participate in regulating sleep duration under different photoperiods (Wood and Loudon, 2014).

To address our hypothesis, we tested fruit flies mutant for different circadian clock genes under a range of photoperiods. We found that flies lacking the circadian photoreceptor CRYPTOCHROME (CRY) display increased sleep duration specifically under short photoperiods, similar to the winter hypersomnolence in SAD. Genetic and pharmacological analysis identified that CRY is functioning in GABAergic neurons and acts upon GABA-A receptor to promote wakefulness. We further narrowed down the neural circuitry mediating the influences of CRY on sleep by demonstrating that *cry* deficiency increases the GABAergic tone and reduces calcium concentration in the wake-promoting large ventral lateral neurons (l-LNvs) which are known to be GABA-A+ (Hamasaka et al., 2005; Parisky et al., 2008). Consistentl with previous data, inhibiting these neurons increases sleep while activating them blocks the effects of *cry* deficiency on sleep. CRY may function in part in the GABAergic small ventral lateral neurons (s-LNvs), and lack of *cry* increases GABA level and the activity of these cells while impairing their GABA transmission suppresses the sleep phenotype of *cry* mutants (Allada and Chung, 2010). In summary, here we identify a potential role for CRY in down-regulating GABAergic signaling specifically under short photoperiod. This in turn enhances the neural activity of the wake-promoting l-LNvs, potentially contributing to the increased wakefulness during short winter-like days. These findings reveal a mechanism underlying how sleep duration is determined under winter-like photoperiod, while disruptions of this regulatory system may be related to the winter hypersomnolence associated with SAD.

## Results

### *cry* mutation increases sleep duration specifically under short photoperiods

We assessed the sleep of flies mutant for circadian clock gene *period* (*per*^0^), *timeless* (*tim^0^*), *clock* (*clk^jrk^*), *cycle* (*cyc^0^*) and *cry* (*cry^b^*) under a range of photoperiods (Figure 1A and B)(Allada, 1998; Konopka and Benzer, 1971; Sehgal et al., 1994; Stanewsky, 1998). We found that *cry^b^* mutation, which is known to be a loss of function or severe hypomorphic allele, leads to increased sleep duration under 4 h light: 20 h dark condition (4L20D) and 8L16D but not under longer photoperiods (Stanewsky, 1998). Since this phenotype recapitulates the winter hypersomnolence of SAD patients, we further characterized the effects of *cry* deficiency on sleep. We focused on sleep under 4L20D, as the extent of sleep increase is slightly larger than that of 8L16D. We also demonstrated that *cry* mutation lengthens sleep duration in both male and female flies, while waking activity is not significantly reduced in the mutants (Figure 1C and D). This means the increased sleep in the mutants is not caused by defects of locomotor ability. We next examined the sleep architecture of these flies and found that *cry* mutation enhances sleep by extending the duration of average sleep bout rather than increasing sleep bout number, indicating that *cry* deficiency promotes sleep consolidation under short photoperiod (Figure 1E and F). Because CRY exerts influences on sleep/wakefulness in a gender-independent manner, we used male flies for the remainder of the study. In addition, we tested the effects of a *cry* knock-out allele (*cry^0^*) on sleep under 4L20D and also observed significantly prolonged sleep duration, similar to *cry^b^* mutation (Figure 1—figure supplement 1) (Dolezelova et al., 2007).

**Figure 1.**
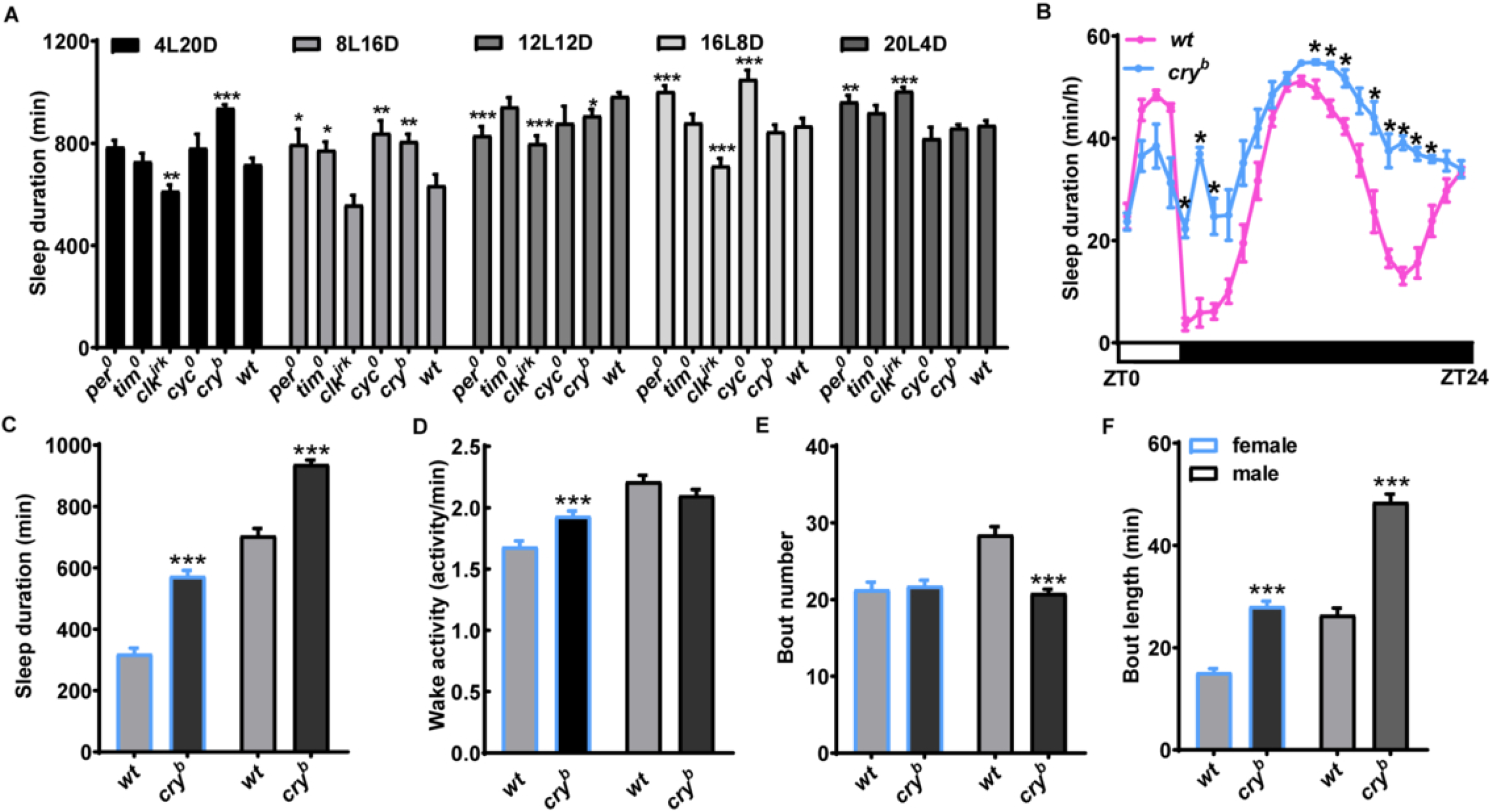
*cry* mutation increases sleep duration selectively under short photoperiods. (A) Daily sleep duration in male circadian clock gene mutants *per*^0^, *tim^0^*, *clk^jrk^*, *cyc^0^*, and *cry^b^* compared to wild-type (WT) males under 4L20D, 8L16D, 12L12D, 16L8D, 20L4D (12L12D, n = 86, 63, 91, 14, 91, 53 flies; 8L16D, n = 21, 31, 31, 10, 32, 31 flies; 12L12D, n = 31, 14, 32, 14, 27, 32 flies; 16L8D, n = 36, 30, 33, 17, 36, 30 flies; 20L4D, n = 42, 55, 83, 26, 91, 89 flies). (B) Average sleep traces of *cry^b^* and WT male flies under 4L20D. (C-F) The daily sleep duration (C), waking activity (D), sleep bout number (E), average sleep bout length (F) for *cry^b^* and WT female and male flies under 4L20D (n = 56, 88, 58, 91 flies). For (B), statistical differences between *cry^b^* and WT is determined by paired two-tailed *t*-test, *\*P* < 0.001. For A and C-E, statistical differences is determined by two-tailed Student’s *t*-test, \*\*\**P* < 0.001. Error bars represent standard error of the mean (SEM). ZT, Zeitgeber Time (ZT0 is the time of lights on).

### CRY functions in GABAergic neurons and promotes wakefulness via GABA/GABA-A

Upon light activation, CRY binds to the clock protein TIM and results in its degradation, which is followed subsequently by degradation of CRY itself (Busza et al., 2004; Ceriani et al., 1999; Lin et al., 2001). Therefore, we first tested whether the influences of CRY on sleep/wakefulness requires TIM. We monitored sleep in flies mutant for both *cry* and *tim*, and found that the sleep duration of these double mutants are comparable to that of *cry* mutants (Figure 2—figure supplement 1). This indicates that CRY promotes wakefulness in a TIM-independent manner.

**Figure 2.**
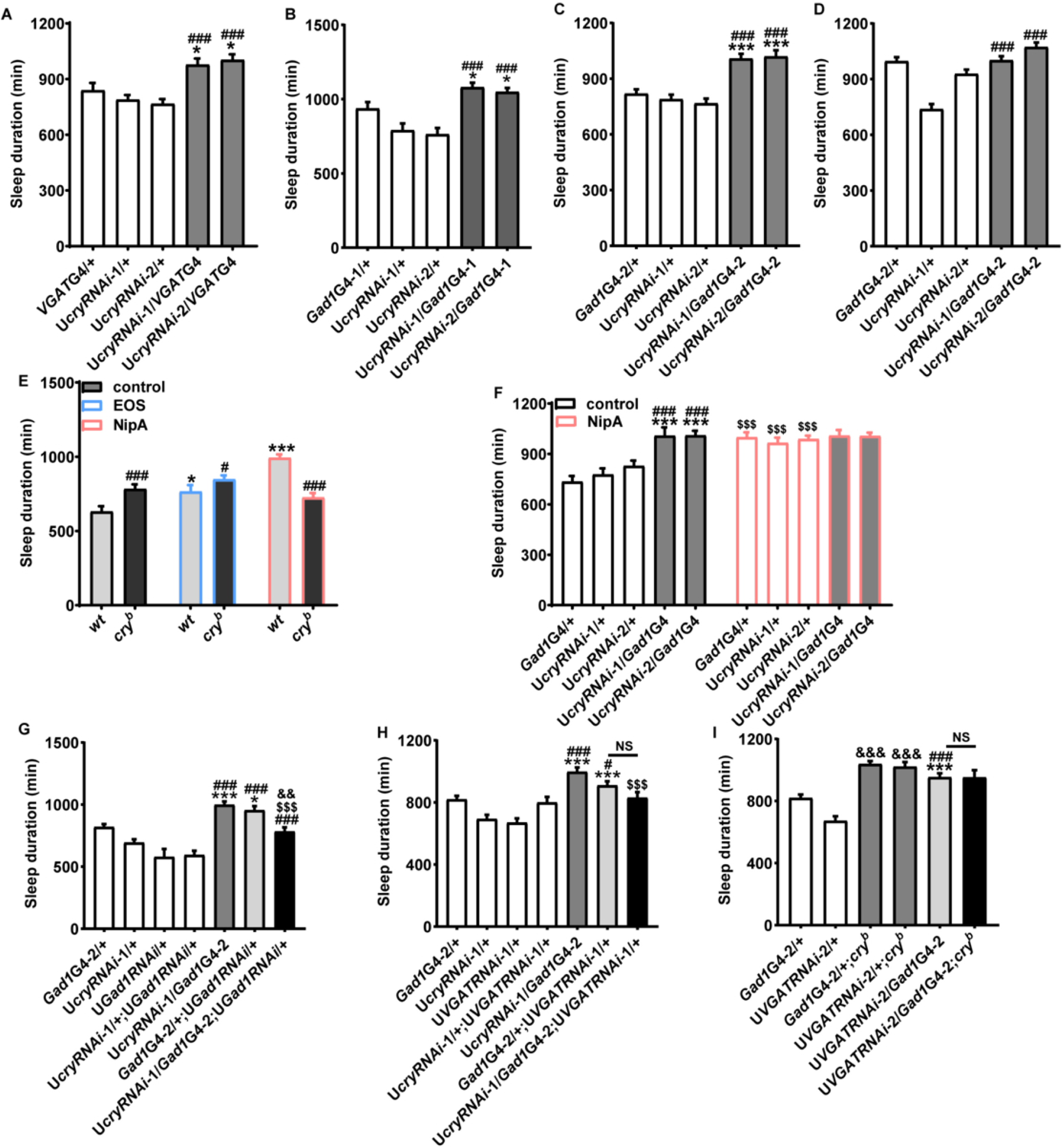
CRY functions in GABAergic neurons and promotes wakefulness via GABA signaling. (A, B, C) Daily sleep duration of male flies with *cry* knocked down in GABAergic neurons by *VGAT*GAL4 (A) (n = 34, 31, 31, 25, 23 flies), *Gad1*GAL4-1 (B) (n = 29, 26, 29, 23, 32 flies) or *Gad1*GAL4-2 (C) (n = 72, 74, 96, 49, 54 flies) monitored under 4L20D. (D) Daily sleep duration of *cry* knocked down in GABAergic neurons by *Gad1*GAL4 monitored under 12L12D (n = 20, 19, 30, 26, 32, 20 flies). For A-D, One-way ANOVA with Bonferroni multiple comparison test: compared to GAL4 control, \*\*\**P* < 0.001; compared to UAS control, #*P* < 0.05, ###*P* < 0.001. (E) Daily sleep duration of male WT and *cry^b^* flies fed with EOS or NipA under 4L20D (n = 31, 32, 30, 30, 25, 32 flies). Two-tailed Student’s *t*-test: compared to WT, ###*P* < 0.001; compared to vehicle control, \**P* < 0.05, \*\*\**P* < 0.001. (F) Daily sleep duration of *cry* RNAi and control flies fed with NipA under 4L20D (n = 29, 19, 26, 15, 28, 23, 24, 26, 15, 24 flies). For comparison between RNAi flies vs. UAS/GAL4 controls, one-way ANOVA with Bonferroni multiple comparison test was used: compared to GAL4 control, \*\*\**P* < 0.001; compared to UAS control, ###*P* < 0.001. For comparing to vehicle control, two-tailed Student’s *t*-test was used, $$$*P* < 0.001. (G) Daily sleep duration of male flies with *cry* and *gad1* knocked down in GABAergic neurons monitored under 4L20D (n = 72, 73, 24, 33, 45, 33, 27 flies). For comparison between RNAi flies vs. UAS/GAL4 controls, one-way ANOVA with Bonferroni multiple comparison test was used: compared to GAL4 control, \*\**P* < 0.01, \*\*\**P* < 0.001; compared to UAS control, ###*P* < 0.001. For comparing to RNAi control, two-tailed Student’s *t*-test was used: compared to UAS*cry*RNAi/GAL4 control, $$$*P* < 0.001; compared to UAS*Gad1*RNAi/GAL4 control, &&&*P* < 0.001. (H) Daily sleep duration of male flies with *cry* and *VGAT* knocked down in GABAergic neurons monitored under 4L20D (n = 72, 73, 46, 41, 45, 66, 34 flies). For comparison between RNAi flies vs. UAS/GAL4 controls, one-way ANOVA with Bonferroni multiple comparison test was used: compared to GAL4 control, \*\*\**P* < 0.001; compared to UAS control, #*P* < 0.05, ### *P* < 0.001. For comparing to RNAi control, two-tailed Student’s *t*-test was used: compared to UAS*cry*RNAi/GAL4 control, $$$*P* < 0.001; compared to UAS*VGAT*RNAi/GAL4 background, not significant. (I) Daily sleep duration of *cry* mutant flies with *VGAT* knocked down in GABAergic neurons monitored under 4L20D (n = 72, 59, 20, 23, 69, 22 flies). For comparison between RNAi flies vs. UAS/GAL4 controls, one-way ANOVA with Bonferroni multiple comparison test was used: compared to GAL4 control, \*\**P* < 0.01, \*\*\**P* < 0.001; compared to UAS control, #*P* < 0.05, ###*P* < 0.001. For comparison between mutant vs. control, two-tailed Student’s *t*-test was used: compared to WT background, &&&*P* < 0.001; compared to U*VGAT*RNAi/GAL4, not significant. Error bars represent SEM; G4, GAL4; U, UAS; NS, not significant.

To identify anatomical substrates that mediate the effects of CRY on wakefulness, we employed the UAS/GAL4 system to knock down *cry* in different brain structures and cell types and verified that *cry* is indeed knocked down by assessing its mRNA level (Figure 2—figure supplement 2). We found that knocking down *cry* in GABAergic neurons using *vesicular GABA transporter* (*VGAT*)GAL4 or two independent *glutamic acid decarboxylase 1* (*Gad1*)GAL4 lines results in increased sleep duration under 4L20D but not 12L12D, similar to the *cry* mutant phenotype (Figure 2A-D; Figure 2—figure supplement 3A-3L)(Deng et al., 2019). Given that *Gad1*GAL4-2 generated a more prominent sleep phenotype, we used this driver for the remainder of this study.

Since CRY appears to act in GABAergic neurons to promote wakefulness, we tested whether GABA is involved in this regulatory process. We first took a pharmacological approach and fed flies with drugs to inhibit GABA transaminase (ethanolamine-O-sulphate, EOS) or GABA transporter (nipecotic acid, NipA). Both of these treatments are known to increase GABA level, and indeed lengthens the sleep duration in WT flies (Ki and Lim, 2019; Leal and Neckameyer, 2002). *cry* mutation fails to enhance sleep when treated with NipA, while the extent of sleep increase caused by *cry* mutation is smaller when treated with EOS compared to the control (Figure 2E; Figure 2—figure supplement 4A-4E). Similarly, knocking down *cry* in GABAergic neurons no longer lengthens sleep duration after NipA treatment (Figure 2F; Figure 2—figure supplement 4F-4I). For further validation, we adopted a genetic approach. We found that knocking down *Gad1*, the synthetic enzyme for GABA, suppresses the long sleep phenotype of *cry* RNAi flies (Figure 2G; Figure 2—figure supplement 5A-5D) (Jackson et al., 1990). In addition, we knocked down *VGAT* which encodes an essential transporter responsible for packing GABA into synaptic vesicles (Enell et al., 2007). *cry* deficiency fails to lengthen sleep duration when *VGAT* is knocked down (Figure 2H and I; Figure 2—figure supplement 5E-5L). We verified that *Gad1* and *VGAT* are indeed knocked down by measuring their mRNA levels (Figure 2—figure supplement 5M and N). As a control for the genetic interaction experiments, we co-expressed a GFP RNAi and found this does not significantly alter the sleep duration of *cry* RNAi flies (Figure 2—figure supplement 5O). These series of results indicate that CRY acts via GABA to promote wakefulness.

We next sought to identify GABA receptor that mediates the effects of CRY on sleep/wakefulness. We fed flies with agonists of GABA-A (THIP) and GABA-B receptor (SKF-97541) (Dissel et al., 2015; Ki and Lim, 2019; Matsuda et al., 1996; Mezler et al., 2001). Both drugs enhance sleep in WT, while *cry* mutation can increase sleep in flies fed with SKF-97541 but not THIP, implicating that CRY acts through GABA-A to promote wake (Figure 3A-C; Figure 3—figure supplement 1A-F). Consistent with previous data, GABA-A receptor antagonist carbamezapine (CBZ) reduces sleep in WT flies while *cry* mutation fails to lengthen sleep duration after CBZ treatment (Figure 3D and E; Figure 3—figure supplement 1G-I) (Agosto et al., 2008). We also treated *cry* RNAi flies with THIP or CBZ and found this treatment abolished the long sleep phenotype as well (Figure 3F-I; Figure 3—figure supplement 2A-F). To validate that CRY modulates sleep/wakefulness by acting upon GABA-A receptor, we tested for genetic interaction between *cry* and *Resistant to dieldrin* (*Rdl*), a gene that encodes GABA-A receptor in flies and has previously been shown to be involved in sleep regulation (Chung et al., 2009; ffrench-Constant et al., 1993; Parisky et al., 2008). We found that *Rdl* mutation (*Rdl^MDRR^*) blocks the sleep-enhancing effect of *cry* RNAi (Figure 3J-L; Figure 3—figure supplement 3)(Agosto et al., 2008). These findings demonstrate that GABA-A receptor mediates the wake-promoting function of CRY.

**Figure 3.**
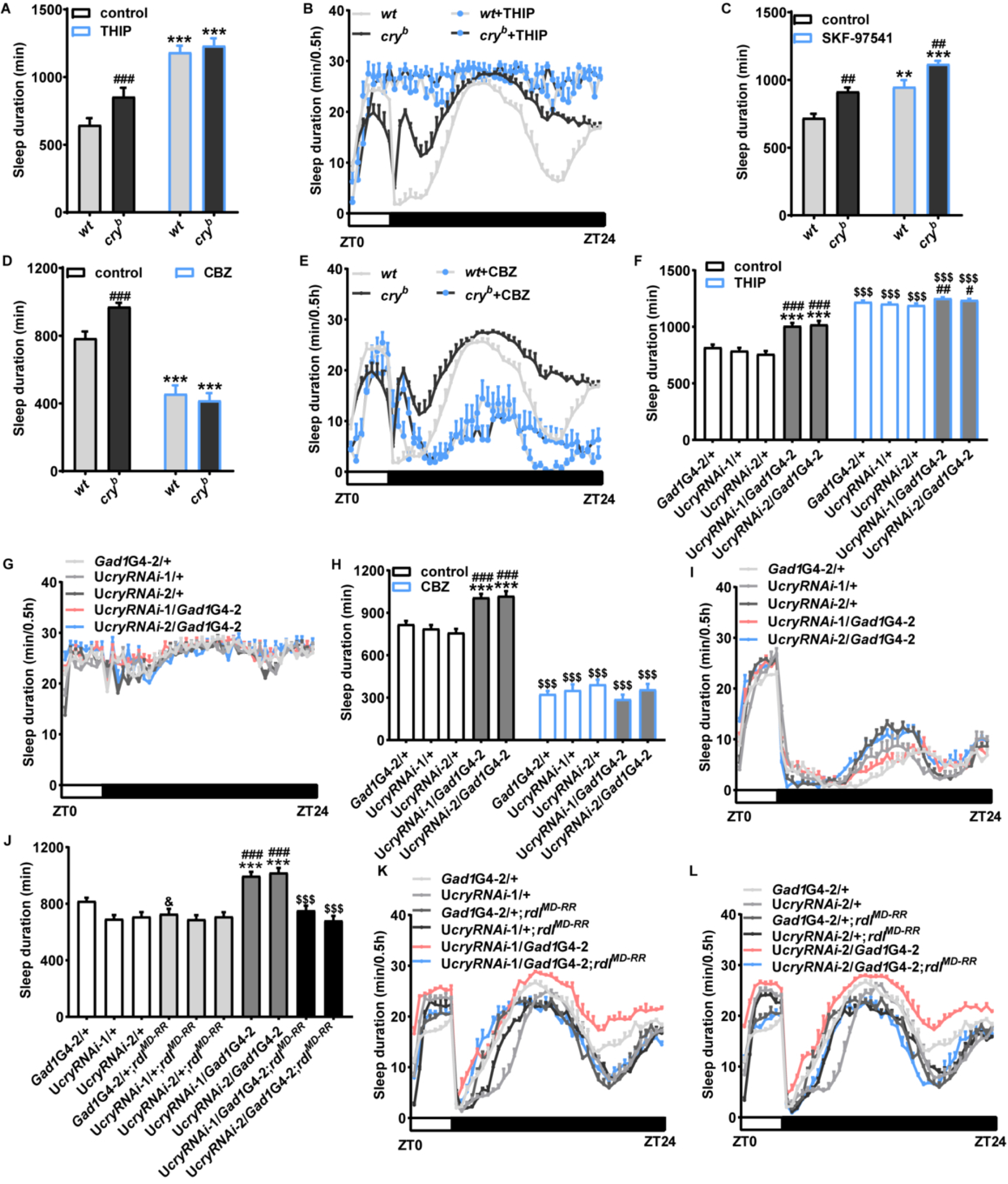
GABA-A receptor mediates the effects of CRY on sleep/wakefulness. (A, C, D) Daily sleep duration of male WT and *cry* mutant flies fed with THIP (A) (n = 28, 29, 16, 19 flies), SKF-97541 (C) (n = 25, 21, 23, 18 flies) or CBZ (D) (n = 28, 25, 31, 29 flies) under 4L20D. Two-tailed Student’s *t*-test: compared to WT, ###*P* < 0.001; compared to vehicle control, \*\*\**P* < 0.001.(B, E) Sleep profile of male WT and *cry* mutant flies fed with THIP (B) or CBZ (E) under 4L20D in A and D, respectively. White box indicates light period while black box indicates dark period. (F, H) Daily sleep duration of male *cry* RNAi and control flies fed with THIP (F) (n = 32, 32, 32, 32, 29, 23, 30, 27,27, 20 flies) or CBZ (H) (n = 24, 16, 18, 22, 23, 18, 22, 26, 30, 24 flies) under 4L20D. For comparison between RNAi flies vs. UAS/GAL4 controls, one-way ANOVA with Bonferroni multiple comparison test was used: compared to GAL4 control, \*\*\**P* < 0.001; compared to UAS control, ##*P* < 0.01, ###*P* < 0.001. For comparing to vehicle control, two-tailed Student’s *t*-test was used, $$$*P* < 0.001. (G, I) Sleep profile of male *cry* RNAi and control flies fed with THIP (G) or CBZ (I) under 4L20D in F and H, respectively. White box indicates light period while black box indicates dark period. (J) Daily sleep duration of male *rdl^MD-RR^*flies with *cry* knocked down in GABAergic neurons under 4L20D, along with relevant controls (n = 72, 73, 66, 44, 45, 33, 45, 54, 38, 33 flies). For comparison between RNAi flies vs. UAS/GAL4 controls, one-way ANOVA with Bonferroni multiple comparison test was used: compared to GAL4 control, \*\*\**P* < 0.001; compared to UAS control, ###*P* < 0.001. For comparing to WT or UAS*cry*RNAi/GAL4 control, two-tailed Student’s *t*-test was used: compared to WT, &*P* < 0.05; compared to UAS*cry*RNAi/GAL4 control, $$$*P* < 0.001. (K, L) Sleep profile of male *rdl^MD-RR^* flies with *cry* knocked down in GABAergic neurons using *cry*RNAi-1 (K) or *cry*RNAi-2 (L) under 4L20D in J. Error bars represent SEM; G4, GAL4; U, UAS.

In short, pharmacological and genetic approaches reveal that CRY acts in GABAergic neurons and impinges on GABA/GABA-A signaling to promote wakefulness.

### CRY may act upon l-LNvs to promote wakefulness

Previous studies reported that GABA enhances sleep at least in part by activating RDL on the l-LNvs, thus inhibiting the activities of these cells and reducing the release of the arousal-promoting neuropeptide pigment dispersing factor (PDF) (Chung et al., 2009; Parisky et al., 2008). Therefore, we tested whether the l-LNvs mediate the effects of CRY on sleep/wakefulness. We first tested whether the l-LNvs indeed receive projections from GABAergic neurons. We expressed a GFP-tagged synaptotagmin (*syt*-*GFP*) which labels axon terminals using *Gad1*GAL4, and GFP signal was observed at the l-LNv soma (Figure 4—figure supplement 1A) (Zhang et al., 2002). We employed the GFP Across Synaptic Partners (GRASP) method with enhanced specificity (t-GRASP) to check for synaptic connections between GABAergic neurons and l-LNvs (Shearin et al., 2018). Pre-mGRASP was expressed in the axon terminal of GABAergic neurons (using *Gad1*GAL4), whereas its t-GRASP partner post-mGRASP was expressed in the l-LNvs (using the driver *Pdf*LexA) (Shang et al., 2008). GFP signal can be detected at the l-LNv soma, which means GABAergic neurons may send synaptic projections to the l-LNvs (Figure 4—figure supplement 1B). We further validated this connection using the *trans*-Tango technique which labels down-stream synaptic targets of GABAergic neurons with the HA tag, and we were able to observe HA expression in the l-LNvs (Figure 4—figure supplement 1C) (Talay et al., 2017). All in all, these results strongly suggest that GABAergic neurons project to the l-LNvs and form synaptic connections. Consistently, knocking down *Rdl* using a R78G01GAL4 line which drives expression in the l-LNvs along with other cells rescues the long sleep phenotype of *cry* mutants (Figure 4—figure supplement 1D) (Yoshii et al., 2015).

**Figure 4.**
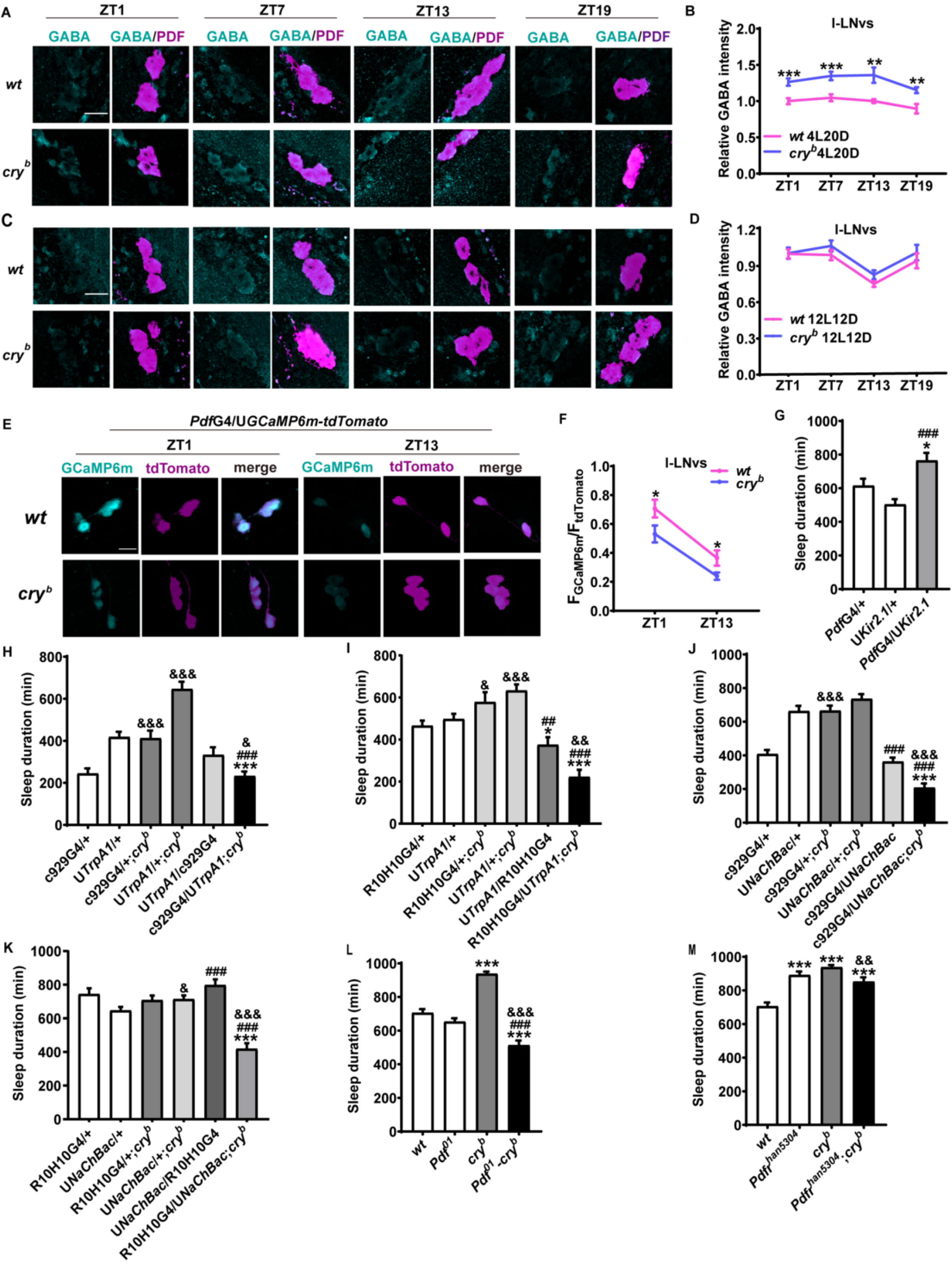
CRY increases the neural activity of the l-LNvs to promote wakefulness. (A-D) Brains from male WT and *cry* mutant flies dissected at ZT1, 7, 13, 19 under 4L20D (A) or 12L12D (C) are immunostained with GABA (cyan) and PDF (magenta) antisera, and representative l-LNvs are displayed. Merged signal is shown as yellow. Bar graphs represent normalized GABA intensity in the l-LNvs under 4L20D (B) (n = 26-56 cells) and 12L12D (D) (n = 32-128 cells). The average value of the control group at ZT1 is set to 1. (E) Representative live image of the l-LNvs expressing GCaMP6m and tdTomato using *Pdf*GAL4.Brain samples are dissected at the indicated time points under 4L20D. (F) Quantification of GCaMP6m signal intensity normalized to that of tdTomato (n = 25-54 cells). Student’s *t*-test: \**P* < 0.05, \*\**P* < 0.01, \*\*\**P* < 0.001. (G) Daily sleep duration of male flies expressing Kir2.1 in PDF neuron using *Pdf*GAL4 and controls, monitored under 4L20D (n = 32, 42, 30 flies). One-way ANOVA with Bonferroni multiple comparison test: compared to GAL4 control, \**P* < 0.05; compared to UAS control, ###*P* < 0.001. (H, I) Daily sleep duration of male *cry* mutant flies expressing TrpA1 in the l-LNvs using c929GAL4 (H) (n = 42, 42, 38, 35, 45, 37 flies) or R10H10GAL4 (I) (n = 62, 62, 17, 30, 29, 30 flies) and relevant controls, monitored under 4L20D and 29℃ to activate TrpA1. For comparison with UAS/GAL4 controls, one-way ANOVA with Bonferroni multiple comparison test was used: compared to GAL4 control, \*\*\**P* < 0.001; compared to UAS control, ###*P* < 0.001. For comparison between mutant vs. control, two-tailed Student’s *t*-test was used: &*P* < 0.05, &&&*P* < 0.001. (J, K) Daily sleep duration of *cry* mutant flies expressing NachBac in the l-LNvs using c929GAL4 (J) (n = 45, 30, 32, 48, 55, 48 flies) or R10H10GAL4 (K) (n = 25, 59, 29, 74, 25, 32 flies) and relevant controls, monitored under 4L20D. For comparison with UAS/GAL4 controls, one-way ANOVA with Bonferroni multiple comparison test was used: compared to GAL4 control, \**P* < 0.05, \*\*\**P* < 0.001; compared to UAS control, ###*P* < 0.001. For comparison between mutant vs. control, two-tailed Student’s *t*-test was used: &*P* < 0.05, &&&*P* < 0.001. (L, M) Daily sleep duration of male *Pdf^01^-cry^b^* (L) (n = 58, 91, 89, 31 flies) and *Pdfr^han5304^*;*cry^b^*(M) (n = 58, 91, 31, 31 flies) mutants along with relevant controls, monitored under 4L20D. Two-tailed Student’s *t*-test: compared to WT background, \*\*\**P* < 0.001; compared to *Pdf^01^*or *Pdfr^han5304^*, ###*P* < 0.001; compared to *cry^b^*, &&*P* < 0.01, &&&*P* < 0.001. The scale bar represents 15 µm. Error bars represent SEM. G4, GAL4; U, UAS.

We next examined the effects of *cry* mutation on GABA level at the l-LNv soma by immunostaining. As expected, GABA signal is significantly increased in *cry* mutants under 4L20D but not 12L12D (Figure 4A-D). Consistent with elevated GABA, GCaMP6m signal, an indicator of intracellular calcium concentration, is significantly reduced in the l-LNvs under 4L20D (Figure 4E and F) (Chen et al., 2013). This implies that the neural activity of these cells is down-regulated, possibly due to increased GABA signaling.

To test whether the long sleep phenotype in *cry* mutants is due to reduced activity of the l-LNvs, we electrically silenced these neurons (along with other PDF+ neurons) by expressing an inward rectifying potassium channel Kir2.1 (Baines et al., 2001). This results in lengthened sleep duration under 4L20D, mimicking the sleep phenotype of *cry* mutants (Figure 4G; Figure 4—figure supplement 2A-D). On the other hand, activating the l-LNvs (along with other cells) by expressing the temperature-gated depolarizing cation channel TrpA1 or the bacterial depolarization-activated sodium channel NachBac blocks the sleep enhancing effect of *cry* mutation (Figure 4H-K; Figure 4—figure supplement 2E-T) (Hamada et al., 2008; Nitabach et al., 2006). These findings support the notion that *cry* mutation may lengthen sleep duration by down-regulating the neural activity of the l-LNvs. Since it has been shown that the l-LNvs promote arousal by releasing PDF, we assessed whether PDF or its receptor PDFR is required by CRY to exert influence on sleep/wakefulness (Chung et al., 2009; Parisky et al., 2008). *cry* mutation fails to lengthen sleep duration on *Pdf* or *Pdfr* mutant background, indicating the necessity of PDF signaling in mediating the arousal-promoting function of CRY (Figure 4L and M; Figure 4—figure supplement 3) (Hyun et al., 2005; Lear et al., 2005; Mertens et al., 2005; Renn et al., 1999).

To summarize, these results suggest that CRY promotes wakefulness by reducing GABA signaling and thus increasing the activity of the l-LNvs, which may in turn lead to increased release of PDF.

### CRY may act in the s-LNvs to promote wakefulness

To identify the subset of GABAergic neurons in which CRY functions to regulate wakefulness, we first examined the expression pattern of *Gad1*GAL4 by labeling the GAL4+ cells with a nuclear GFP (nls-GFP) (Shiga et al., 1996). While the l-LNvs are not GFP+, we noticed that the s-LNvs which are close to the l-LNvs and also express PDF appear to be GFP+ (Figure 5A and B; Figure 5-figure supplement 1A). In addition, nls-GFP was expressed in GABAergic neurons (using *Gad1*GAL4), whereas RFP was expressed in the l– and s-LNvs (using the driver *Pdf*LexA). GFP signal can be detected in the s-LNvs but not the l-LNvs (Figure 5-figure supplement 1B). Because the s-LNvs are known to express CRY, we suspected that CRY may be acting in these s-LNvs to regulate the activity of the l-LNvs via GABA signaling (Benito et al., 2008; Yoshii et al., 2008). To test this idea, we knocked down *cry* using R6GAL4 (which drives expression in the s-LNvs and several other cells in the brain) while over-expressing *dicer2* (*dcr2*) to enhance RNAi efficiency and observed a modest but significant lengthening of sleep duration (Figure 5C; Figure 5—figure supplement 2A-D) (Helfrich-Forster et al., 2007). However, when we adopted a *Pdf*GAL80 to block the actions of GAL4 in the PDF neurons in *Gad1*GAL4/UAS*cry*RNAi, this does not alter the long-sleep phenotype (Figure 5—figure supplement 2E) (Stoleru et al., 2004). These series of results suggest that *cry* expression in the s-LNvs may be necessary but not sufficient to maintain normal sleep/wakefulness. We were indeed able to detect GABA signal at the s-LNvs, while GABA intensity is enhanced in *cry* mutants under 4L20D but not 12L12D (Figure 5D-G). To further validate that the s-LNvs are GABAergic, we knocked down *VGAT* in these cells and observed a decrease of GABA intensity (Figure 5—figure supplement 3A and B).

**Figure 5.**
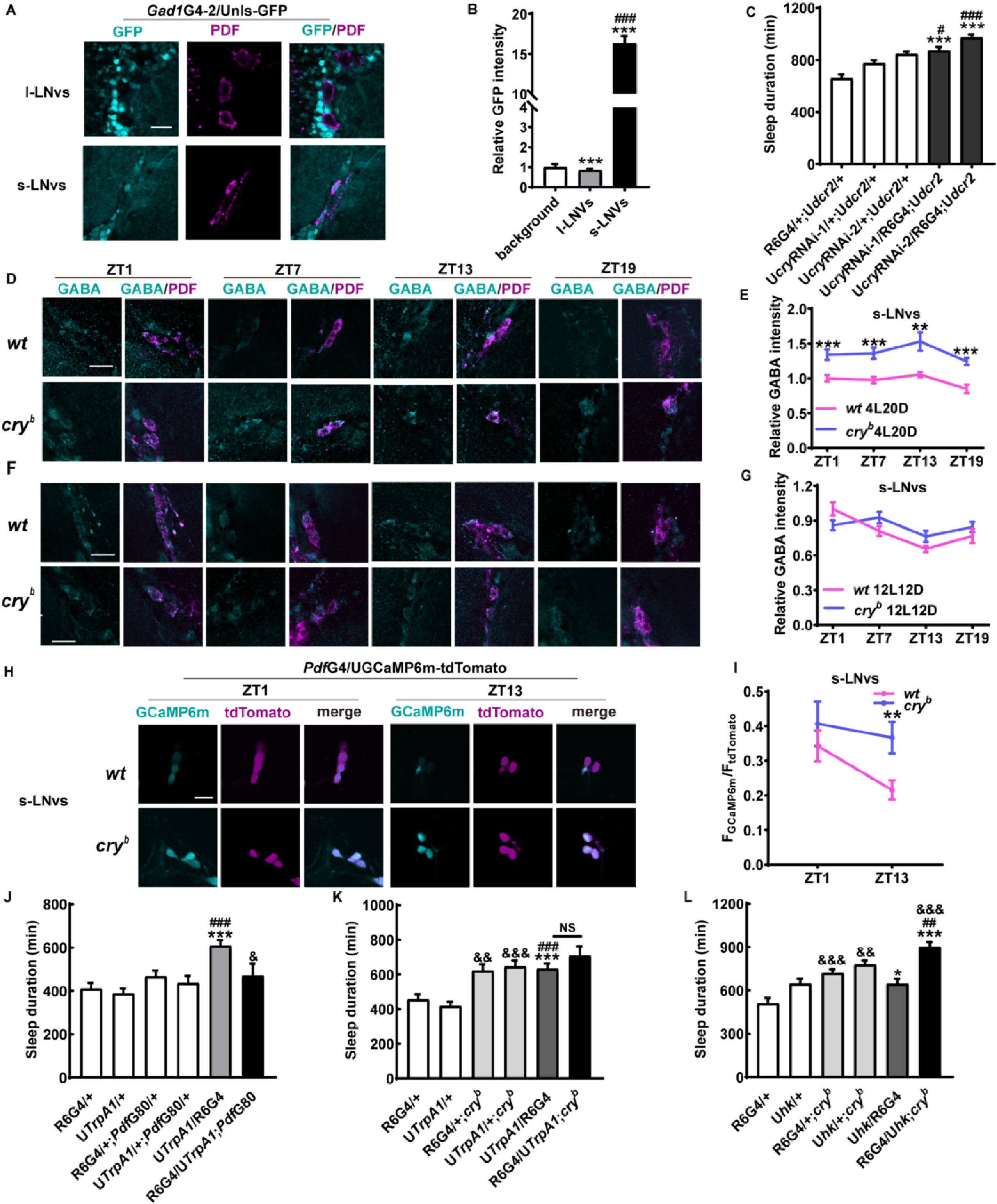
CRY acts in the s-LNvs to inhibit their neural activity and promote wakefulness. (A) Brains of male flies expressing nls-GFP driven by *Gad1*GAL4 maintained under 4L20D and immunostained with PDF antisera. Representative l-LNv and s-LNv are displayed. (B) Quantification of GFP signal intensity of PDF neuron (n = 20, 49, 43 cells). Two-tailed Student’s *t* test: \*\*\**P* < 0.001. (C) Daily sleep duration of flies with *cry* knocked down in the s-LNvs using R6GAL4 and relevant controls, monitored under 4L20D (n = 45, 60, 63, 34, 39 flies). (D-G) Brains from male WT and *cry* mutant flies dissected at ZT1, 7, 13, 19 under 4L20D (D) or 12L12D (F) are immunostained with GABA (cyan) and PDF (magenta) antisera, and representative s-LNvs are displayed. Merged signal is shown as yellow. Bar graphs represent normalized GABA intensity in the s-LNvs under 4L20D (E) (n = 20-53 cells) and 12L12D (G) (n = 29-67 cells). The average value of the control group at ZT1 is set to 1. (H) Representative live image of the s-LNvs expressing GCaMP6m and tdTomato using *Pdf*GAL4. Brain samples are dissected at the indicated time points under 4L20D. (I) Quantification of GCaMP6m signal intensity normalized to that of tdTomato (n = 29-47 cells). Two-tailed Student’s *t*-test: \**P* < 0.05, \*\**P* < 0.01, \*\*\**P* < 0.001. (J) Daily sleep duration of male flies expressing TrpA1 in the s-LNvs using R6GAL4 and relevant controls, monitored under 4L20D and 29℃ to activate TrpA1 (n = 45, 54, 29, 22, 56, 22 flies). (K) Daily sleep duration of male *cry* mutant flies expressing TrpA1 in the s-LNvs using R6GAL4 and relevant controls, monitored under 4L20D and 29℃ to activate TrpA1 (n = 33, 42, 23, 35, 36, 20 flies). (L) Daily sleep duration of male *cry* mutant flies over-expressing HK in the s-LNvs using R6GAL4 and relevant controls, monitored under 4L20D (n = 24, 29, 23, 30, 37, 22 flies). For comparison with UAS/GAL4 controls, one-way ANOVA with Bonferroni multiple comparison test was used: compared to GAL4 control, \**P* < 0.05, \*\*\**P* < 0.001; compared to UAS control, ##*P* < 0.01, ###*P* < 0.001. For comparison between mutant vs. control, two-tailed Student’s *t*-test was used: compared to WT background, &&*P* < 0.01, &&&*P* < 0.001. The scale bar represents 15 µm. Error bars represent SEM. G4, GAL4; U, UAS; NS, not significant.

Next, we examined the effects of *cry* deficiency on the activity level of the s-LNvs. We found that *cry* mutation increases calcium concentration in these cells under 4L20D, in stark contrast to that of the l-LNvs (Figure 5H and I). This strongly suggests that *cry* deficiency leads to elevated neural activity in the s-LNvs. Consistently, we observed increased sleep when we activated these cells using TrpA1, similar to the long-sleep phenotype of *cry* mutants (Figure 5J; Figure 5—figure supplement 4A-D). Because R6GAL4 is also expressed in several other cells in the brain, we combined *Pdf*GAL80 to verify that the sleep-enhancing effect is caused by over-activation of the s-LNvs (Helfrich-Forster et al., 2007). Indeed, we no longer observed the long-sleep phenotype when GAL4 expression is inhibited in the PDF neurons by GAL80 (Figure 5J; Figure 5—figure supplement 4A-D). Moreover, we found that *cry* mutation can no longer exert effects on sleep when the s-LNvs (along with other cells) are activated, further suggesting that the s-LNvs mediate the influences of CRY on sleep (Figure 5K; Figure 5—figure supplement 4E-H). Because CRY has been shown to regulate membrane depolarization and input resistance together with the redox sensor of the voltage-gated potassium channel β-subunit HYPERKINETIC (HK), we tested whether CRY and HK also function together in the s-LNvs to regulate arousal (Agrawal et al., 2017; Fogle et al., 2015). We found that while over-expressing *hk* with R6GAL4 in WT flies does not alter sleep duration under 4L20D, over-expressing it in *cry* mutants significantly increases sleep (Figure 5L; Figure 5—figure supplement 4I-L). This genetic interaction indicates that CRY and HK cooperate to regulate wakefulness, possibly by modulating the electric activity of the s-LNvs.

Next, we expressed syt-GFP using R6GAL4 and observed GFP signal at the soma of the l-LNvs, suggesting that the s-LNvs send axonal terminals to the l-LNvs (Figure 6A). *trans-*Tango technique further implies that the s-LNvs project to form synaptic connections with the l-LNvs (Figure 6B). Moreover, when we disrupted GABA transmission using R6GAL4 by knocking down *VGAT* or *Gad1*, this reduced GABA intensity in the l-LNvs, implicating that the s-LNvs may release GABA onto the l-LNvs (Figure 6C-E). At the behavioral level, these manipulations suppress the long-sleep phenotype of *cry* mutation (Figure 6F-I; Figure 6—figure supplement 1).

**Figure 6.**
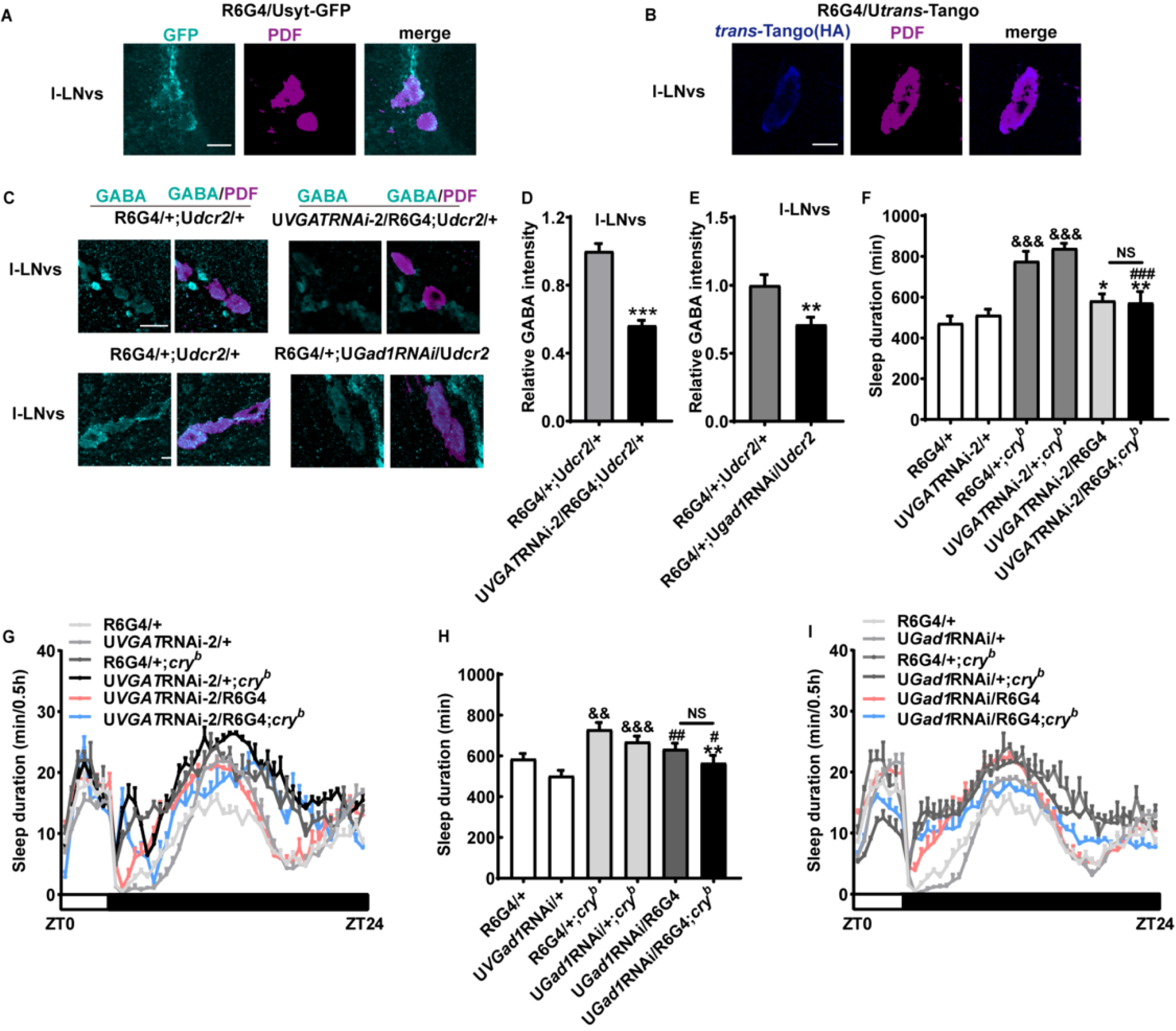
s-LNvs release GABA onto the l-LNvs. (A) l-LNvs of male flies expressing syt-GFP in the s-LNvs using R6GAL4 maintained under 4L20D and immunostained with PDF antisera. (B) l-LNvs of male flies expressing trans-Tango in the s-LNvs using R6GAL4 maintained under 4L20D and immunostained with HA and PDF antisera. (C) Brains from male flies with *VGAT* (top) or *Gad1* (bottom) knocked down in the s-LNvs using R6GAL4 and controls maintained under 4L20D and immunostained with GABA (cyan) and PDF (magenta) antisera. Representative l-LNvs are displayed. Merged signal is shown as yellow. (D, E) Bar graphs represent normalized GABA intensity in the l-LNvs of flies with *VGAT* (D) (n = 29, 37 cells) or *Gad1* (E) (n = 37, 51 cells) knocked down in the s-LNvs using R6GAL4, monitored under 4L20D. The average value of the control group is set to 1. Two-tailed Student’s *t*-test: \*\**P* < 0.01, \*\*\**P* < 0.001. (F) Daily sleep duration of male *cry* mutant flies with *VGAT* knocked down in the s-LNvs using R6GAL4 and relevant controls, monitored under 4L20D (n = 31, 31, 32, 29, 43, 19 flies). For comparison between RNAi flies vs. UAS/GAL4 controls, one-way ANOVA with Bonferroni multiple comparison test was used: compared to GAL4 control, \**P* < 0.05, \*\**P* < 0.01; compared to UAS control, ###*P* < 0.01. For comparison between mutant vs. control, two-tailed Student’s *t*-test was used: compared to WT background, &&&P < 0.001. (G) Sleep profile of male *cry* mutant flies with *VGAT* knocked down in the s-LNvs using R6GAL4 and relevant controls in F, monitored under 4L20D. (H) Daily sleep duration of male *cry* mutant flies with *Gad1* knocked down in the s-LNvs using R6GAL4 and relevant controls, monitored under 4L20D (n = 59, 59, 53, 60, 62, 45 flies). For comparison between RNAi flies vs. UAS/GAL4 controls, one-way ANOVA with Bonferroni multiple comparison test was used: compared to GAL4 control, \**P* < 0.05, \*\**P* < 0.01, \*\*\**P* < 0.001; compared to UAS control, #*P* < 0.05, ##*P* < 0.01. For comparison between mutant vs. control, two-tailed Student’s *t*-test was used: compared to WT background, &&&*P* < 0.001. (I) Sleep profile of male *cry* mutant flies with *Gad1* knocked down in the s-LNvs using R6GAL4 and relevant controls in H, monitored under 4L20D.The scale bar represents 15 µm. Error bars represent SEM. G4, GAL4; U, UAS; NS, not significant.

Taken together, these results suggest that *cry* deficiency in the s-LNvs results in increased neuronal activity and thus enhanced GABA release at the l-LNvs via direct synaptic projections, ultimately leading to decreased wakefulness and increased sleep.

### Short photoperiod reduces GABA level at the l-LNvs

Our findings thus far point to an inhibitory role of CRY on the GABAergic tone, and this in turn removes the inhibition on the neural activities of the l-LNvs. Since CRY is degraded by light, we hypothesized that CRY should exert a stronger effect on GABA under short photoperiod (Emery et al., 1998). Consistently, GABA intensity does appear to be down-regulated at the l-LNvs under 4L20D vs. 12L12D (Figure 7A and B). On the other hand, GABA intensity does not exhibit photoperiod-dependent alteration in the s-LNvs (Figure 7C and D). Presumably, this reduced GABA level at the l-LNvs under 4L20D can result in elevated activation of these cells and increased wakefulness. In line with this, WT flies display shortened sleep duration under 4L20D compared to 12L12D (Figure 7E and F). This sleep reduction is a result of decreased sleep bout length but not bout number, while wake activity is not altered by photoperiod (Figure 7G-I). These observations indicate that short photoperiod hampers sleep maintenance, similar to the effects of CRY.

**Figure 7.**
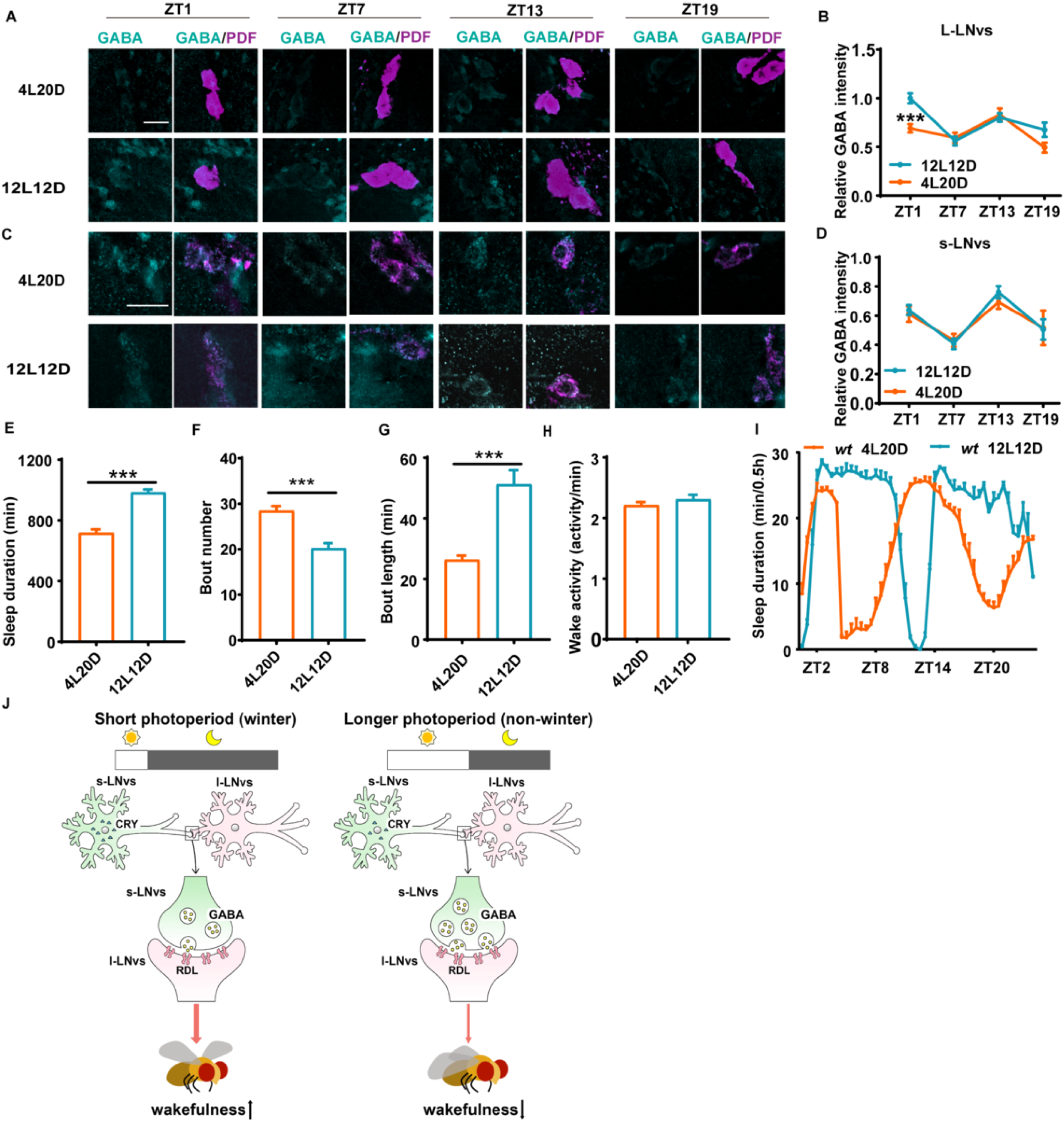
Short photoperiod reduces GABA level at the l-LNvs and increases sleep duration. (A-D) Brains from male WT flies dissected at ZT1, 7, 13, 19 under 4L20D or 12L12D are immunostained with GABA (cyan) and PDF (magenta) antisera. Representative l-LNvs (A) and s-LNvs (C) are displayed. Merged signal is shown as yellow. Bar graphs represent normalized GABA intensity in the l-LNvs (B) (n = 28-55 cells) and s-LNvs (D) (n = 21-59 cells). The average value of the control group at ZT1 is set to 1. Student’s *t-*test: \*\*\**P* < 0.001. (E-I) The daily sleep duration (E), sleep bout number (F), average sleep bout length (G), waking activity (H) and sleep profile (I) of male WT flies under 4L20D and 12L12D (n = 91, 32 flies). Two-tailed Student’s *t*-test: \*\*\**P* < 0.001. The scale bar represents 15 µm. Error bars represent SEM. (J) A model demonstrating how CRY regulates the GABAergic s-LNv/l-LNv circuitry under short vs. longer photoperiods to promote arousal.

## Discussion

Previous studies have shown that CRY mediates light-induced electrical activity of the l-LNvs and acute arousal, which indicates that CRY can immediately act to promote wake and terminate sleep in response to light pulse (Fogle et al., 2015; Sheeba et al., 2008). At the molecular level, this is believed to be accomplished by a direct coupling of light-activated CRY with HK in the l-LNvs (Fogle et al., 2015). Here we found that CRY promotes extended wakefulness under short photoperiod by functioning as an inhibitor of GABAergic tone. In contrast to the previously characterized roles of CRY which are activated by light, we believe this novel function we identify here reflects a role for CRY in the dark. Under short photoperiods, CRY acts in the dark to inhibit GABA signaling and thus promote wakefulness, leading to reduced sleep during the dark phase compared to longer photoperiods. Indeed, CRY is an ideal signal for conveying photoperiodic information, for it is degraded by light and only accumulates during darkness (Emery et al., 1998). It can measure the length of the day/night and thus instruct downstream signaling components to modulate photoperiod-dependent processes. Consistent with this idea, the largest increases of sleep in *cry* mutants occur immediately after lights-off and prior to lights-on (Fig 1B), indicating that CRY functions to promote wakefulness during these time windows which are exactly the time windows that would be affected by photoperiod changes (i.e. there will be light during these time windows under longer photoperiods). In other words, CRY appears to act at the time of the day that is most sensitive to photoperiodic changes, which fits perfectly with the idea that it serves as an instructive signal of day/night length.

While EOS and NipA treatment lengthen sleep duration as previously reported, we are somewhat surprised by the observation that knocking down *Gad1* or *VGAT* in GABAergic neurons also extends sleep duration (Ki and Lim, 2019; Leal and Neckameyer, 2002). We reason this may be due to some sort of over-compensation induced by chronic GABA deficiency to maintain excitation/inhibition balance, as previous studies have reported that the amplitude of glutamatergic current is substantially down-regulated in *Gad1* mutant flies and increased in *Gad1* over-expression flies (Featherstone et al., 2000), (Featherstone et al., 2002). Consistent with this idea, we noticed when *cry* is knocked down in *Gad1/VGAT* RNAi flies, sleep duration is shortened and comparable to that of the controls. This is probably because *cry* depletion enhances the GABAergic tone, thus normalizing GABA signaling in these flies which results in normal sleep duration.

We acknowledge that THIP treatment leads to prominent lengthening of sleep duration, and thus it is possible in this case *cry* deficiency no longer increases sleep duration due to a ceiling effect rather than epistatic interaction. Nonetheless, considering that CBZ and *rdl* mutation also block the effect of *cry* deficiency on sleep duration, it is highly likely that GABA-A receptor mediates the influences of CRY on sleep. One caveat is that the *rdl^MD-RR^* mutation has been shown to diminish the desensitization of GABA-A receptor and is thus believed to be a gain-of-function allele, but here we found that it eliminates the effect of *cry* RNAi on sleep (Zhang et al., 1994). We suspect that similar to *Gad1/VGAT* RNAi, chronic enhancement of GABA-A function associated with *rdl^MD-RR^* mutation may also trigger some kind of compensatory mechanism that counteracts this increased GABA-A activity. Consequently, the influences of *cry* RNAi on sleep are blocked. Of course, a lot more in depth characterizations are required to elucidate these issues.

The s-LNvs have been reported to receive GABAergic inputs possibly via GABA-B receptors, but have not been shown to be able to synthesize GABA (Dahdal et al., 2010; Hamasaka et al., 2005). One study using cell-type specific gene expression profiling demonstrates *Gad1* and *VGAT* expression in both s-LNvs and l-LNvs, although with relatively low signal (Nagoshi et al., 2010). Here we observed that *Gad1*GAL4 is expressed in the s-LNvs, and their GABA intensity is reduced when we use R6GAL4 to knock down *VGAT* in these cells. R6GAL4 drives prominent expression in the s-LNvs with very little if any expression in the l-LNvs, and weaker (and not very consistent) expression in several other neurons in the protocerebrum, pars intercerebralis and subesophageal area which are all believed to lie outside of the circadian neuron network (Helfrich-Forster et al., 2007). Therefore, we reason that the alteration of GABA signal associated with knocking down *VGAT* should arise from VGAT deficiency within the s-LNvs. We do acknowledge that the GABA immunostaining shown here is not optimal, but in combination with the genetic data, the results converge at the conclusion that the s-LNvs are GABAergic as the most plausible explanation. Although knocking down *VGAT* or *Gad1* using R6GAL4 can suppress the long sleep phenotype of *cry* mutants, knocking down *cry* only leads to a modest lengthening of sleep duration. Moreover, inhibiting *cry* RNAi expression in PDF neurons does not eliminate the long-sleep phenotype of *Gad1*GAL4/UAS*cry*RNAi flies. Therefore, we suspect that *cry* deficiency in other GABAergic neurons are also required for the long-sleep phenotype. Given that the s-LNvs are known to express CRY and appear to be GABAergic based on our findings here, we believe that CRY acts at least in part in the s-LNvs to promote wakefulness under short photoperiod.

The molecular mechanism by which CRY down-regulates the GABAergic tone remains unclear. Besides conducting light-driven depolarization via HK, CRY has also been shown to act in synergy with HK to prevent membrane input resistance from falling to a low level in larval salivary glands (Agrawal et al., 2017; Fogle et al., 2015). In contrast to previous studies, here we found that lack of CRY increases the activity of the s-LNvs. Instead of functioning in synergy with HK, CRY appears to act in the opposite direction of HK as over-expressing *hk* enhances the long-sleep phenotype caused by *cry* mutation. We reason that the coupling between CRY and HK as well as their influences on the electric activity in the s-LNvs may be different from that of the l-LNvs and larval salivary glands. Nonetheless, our results also support an interaction between CRY and HK to promote arousal. We suspect that CRY acts via HK to inhibit the activity of the s-LNvs, which results in decreased GABA release and dis-inhibition of the l-LNvs. Extensive further investigations will be needed to elucidate the mechanism by which CRY regulates the activity of the s-LNvs.

In conclusion, here we describe a CRY-controlled GABAergic circuitry potentially involving the l-LNvs and the s-LNvs that adjusts sleep duration in adaptation to changes in day length and propose a mechanistic explanation regarding how this circuitry functions (Fig 7J). Under short photoperiods, more CRY accumulates and inhibits the activity of the GABAergic s-LNvs, leading to a dis-inhibition of the l-LNvs which can release more PDF and promote arousal. Under longer photoperiods, on the other hand, less CRY can accumulate and thus the s-LNvs will exert more inhibitory influences on the l-LNvs, leading to decreased release of PDF and wakefulness. Notably, almost all neurons in the mammalian pacemaker, the suprachiasmatic nucleus, are GABAergic, and GABA/GABA-A signaling have been shown to mediate neuronal coupling in response to photoperiod changes (Ono et al., 2021). We believe a similar GABAergic circuitry may exist in the mammalian system that adjusts sleep/wakefulness to photoperiodic changes.

## Materials and Methods

### Key resource table

Please see Appendix 1.

### Fly strains

All strains were obtained from Bloomington *Drosophila* Stock Center, Vienna Drosophila Resource Center and TsingHua Fly Center or as gifts from colleagues. Except for *Gad1*GAL4-2, all neurotransmitter related GAL4 lines were generated in Dr. Yi Rao’s laboratory (Deng et al., 2019). The Drosophila strains used are listed in the resource table. All flies used for sleep monitoring were backcrossed with the isogenic *w^1118^* strain for at least 5 times except for *cry^03^* which was backcrossed three times. All experiments were conducted in male flies unless otherwise specified.

### Fly sleep monitoring and analysis

Flies were raised on standard cornmeal-yeast-sucrose medium and kept in 12L12D at 25°C until behavior monitoring. ∼3 to 4-day-old flies were entrained under different photoperiods at 25℃ for 4 days, and then their activities in the next 3 days were analyzed. Sleep is defined as 5 min consecutive inactivity. Sleep was analyzed with Counting Macro written in Excel (Microsoft) following previously published protocol (Pfeiffenberger et al., 2010). Flies were fed with agar-sucrose food (2% agar, 5% sucrose) during entire sleep monitoring. TrpA1 flies were raised at 21°C and baseline sleep was monitored at 21°C. Temperature was then raised at lights on to 29°C for further sleep monitoring.

### Drug treatment

For pharmacological experiments, drugs were mixed in the fly food at the following concentrations. For nipecotic acid (10 mg/ml, Sigma) and EOS (10 mM, Sigma), drugs were fed during the entire sleep monitoring. For THIP (10 ug/ml, Sigma), SKF-97541 (10 ug/ml, Tocris) and CBZ (0.15 mg/ml, Sinopharm Chemical Reagent), drugs were fed for 1 day after baseline sleep monitoring. The same amount of solvent was added into the fly food as vehicle control.

### RNA extraction and quantitative real-time PCR (qRT-PCR)

Approximately 50 5-day-old flies were collected and frozen immediately on dry ice. Fly heads were isolated and homogenized in Trizol reagent (Life Technologies). Total RNA was extracted and qRT-PCR conducted following our previously published procedures (Bu et al., 2019).

### Immunostaining

Male flies were entrained for 3 days under indicated photoperiod and collected on Day 4. flies were anesthetized with CO2 and dissected and fixed with 4% paraformaldehyde diluted in PBS at the indicated time points. The brains were then fixed with 4% paraformaldehyde for 30 minutes. Samples were washed with PBT (PBS with 0.3% TritonX-100) for 3 × 10min. Then samples were incubated in PBS with 1% TritonX-100 for 20 min at room temperature. Brain samples were then blocked in PBT with 5% fetal bovine serum (Hyclone) for 30 minutes and subsequently incubated with mouse anti-PDF (1:100, DSHB), rabbit anti-GABA (1:200, Sigma), and anti-HA (1:100, DSHB) for 2-4 days at 4℃ (4 days for anti-GABA and 2 days for the other antibodies). After PBT rinses for 3 times, the brains were incubated with donkey anti-mouse Alexa Fluor-594 (1:1000, Life Technologies), donkey anti-rabbit Alexa Fluor-488 (1:1000, Abcam) and donkey anti-rabbit Alexa Fluor-647 (1:1000, Abcam) overnight at 4℃. Then the brains were rinsed 3 times in PBS and mounted and imaged using Olympus FV3000 confocal microscope with a 60× objective lens. The intensity of GABA signal was quantified by ImageJ software. For each cell, the image slice with the strongest signal in the Z stack was selected and average intensity was quantified. A region on the same image slice was then selected as background and its intensity value was subtracted from the average intensity value of the cell. This subtracted value was subsequently normalized to the average intensity of the control group as indicated in figure legends. Sample sizes in the legends indicate number of cells examined per genotype. For *trans*-Tango experiment, offsprings were raised at 18°C for 3-4 weeks before HA and PDF immunostaining.

### Calcium imaging

For live imaging experiments (GCaMP and tdTomato), flies 2-3 days old were collected into tubes with standard food and entrained under LD for 3 days. Flies were anesthetized with CO2 and brains were dissected at the indicated time points in *Drosophila* adult hemolymph-like saline solution. The dissected brain samples were put on glass slide and sealed with cover slide. The duration for dissection and microscopy should be completed within 0.5 hours for each time point. Images were captured with Olympus FV3000 confocal microscopy with a 20× objective lens. The intensity of GCaMP and tdTomato signals were quantified by ImageJ software. Sample sizes in the legends indicate number of cells examined per genotype.

### Statistical Analysis

For data that fit normal distribution, two-tailed Student’s *t*-test (Microsoft Excel) was used to compare the difference between two genotypes. For data that do not fit normal distribution, Mann-Whitney test (Prism Graphpad) was used to compare the difference between two genotypes. For multiple comparisons, one-way ANOVA with Bonferroni multiple comparison test (Prism Graphpad) was used. Sample size, statistical test, and significance values are indicated in figure legends. Sample size is determined based on previous studies with similar experimental assays.

## Acknowledgements

We would like to thank Drs. Chang Liu, Yi Rao, Liming Wang, Junhai Han, Zhihua Liu and Yi Zhong for kindly providing flies used in this study. We would also like to thank Dr. Chang Liu for helpful advice on GABA immunostaining. This work was supported by grants from the Ministry of Science and Technology of China STI 2030-Major Projects (2021ZD0203202 and 2024YFA1803201), Natural Science Foundation of China (82530044 and 32341021) and Science and Technology Department of Hubei Province (2022CFA049) to Luoying Zhang and grant from the Natural Science Foundation of China (32300984) to Lixia Chen.

## Author contributions

L.C. and L.Z. designed the experiments. L.C. and D.T. conducted the experiments. L.C., C.S. and L.Z. wrote the paper.

## Data Availability

This study includes no data deposited in external repositories.

## Appendix 1

**Appendix 1.**
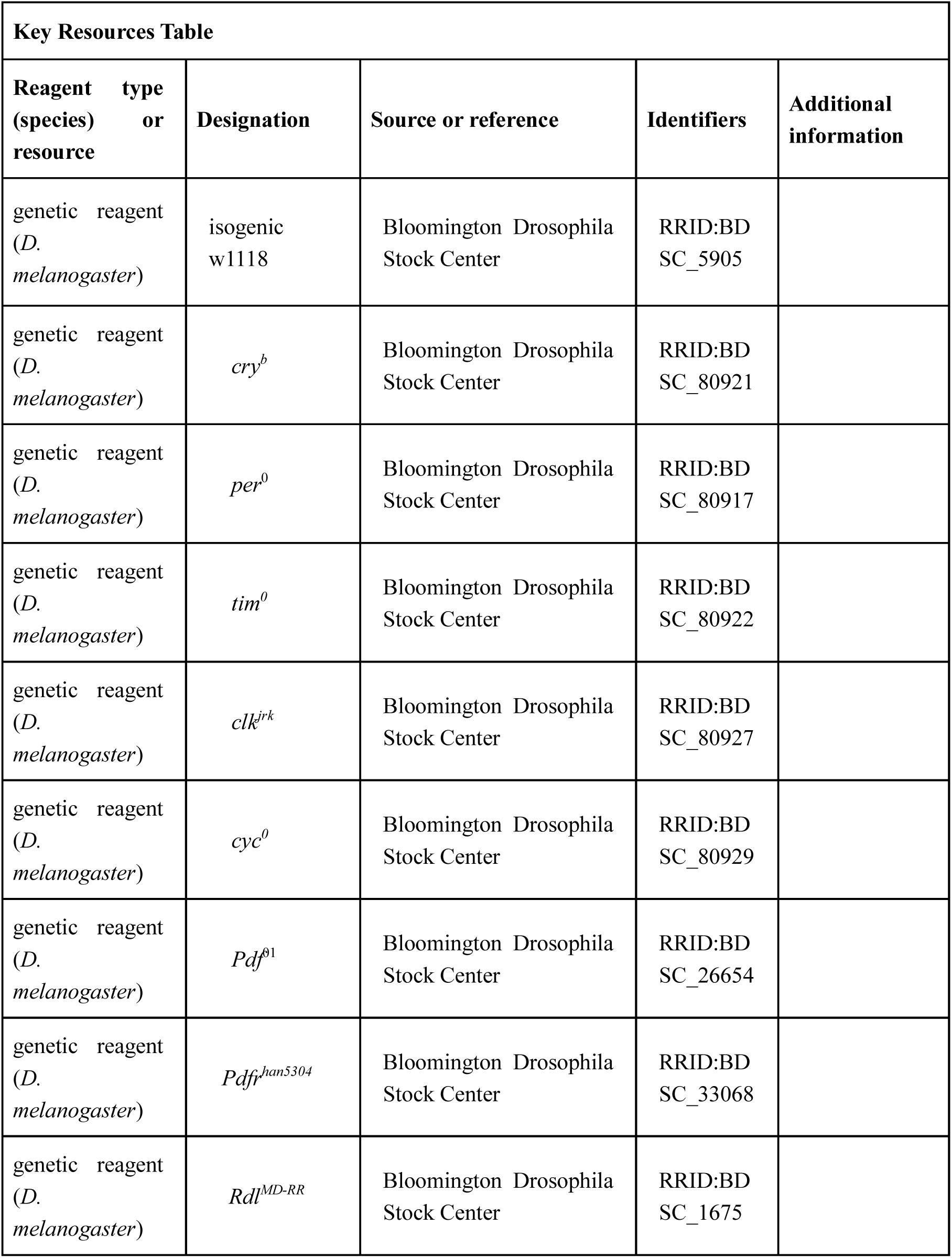
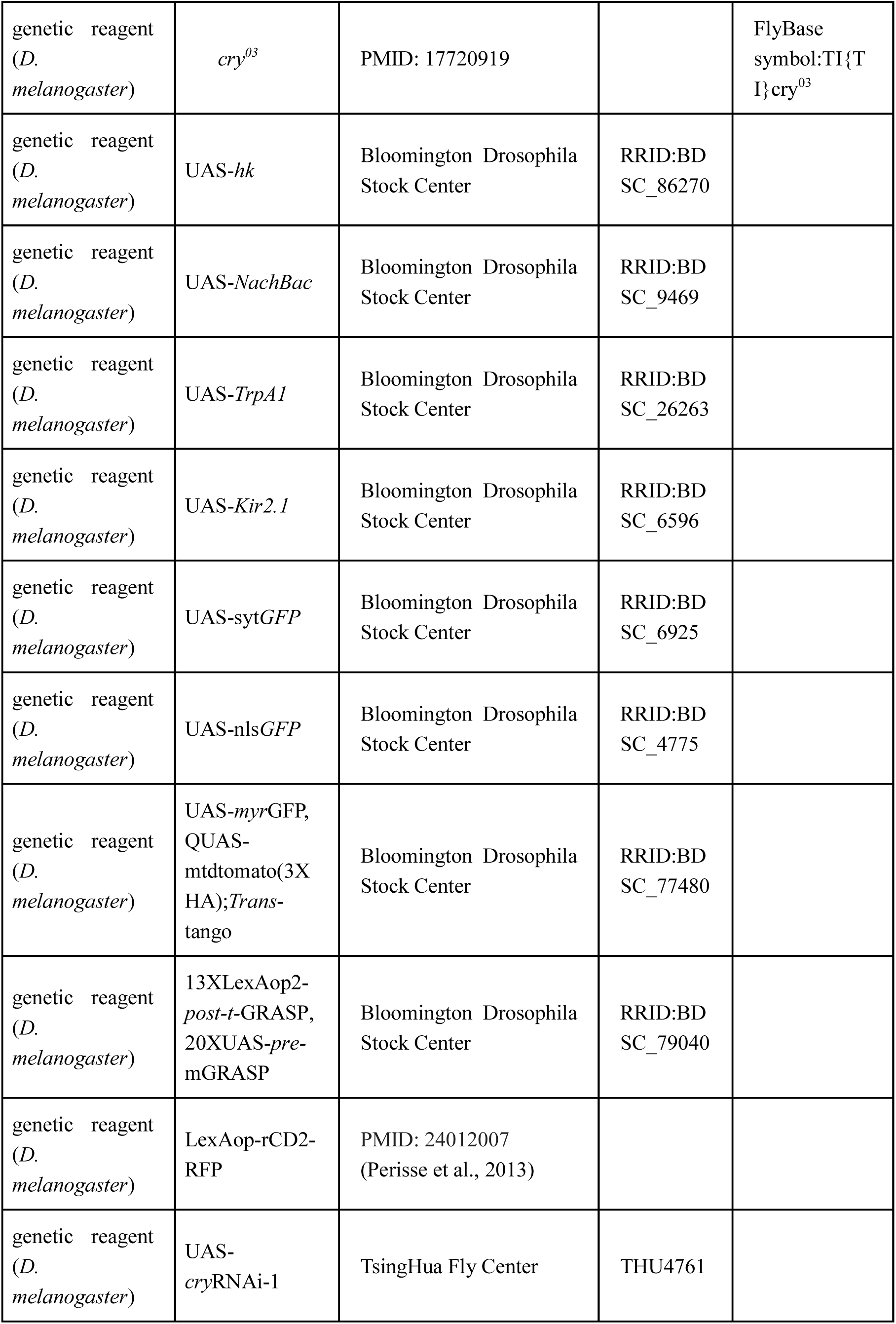
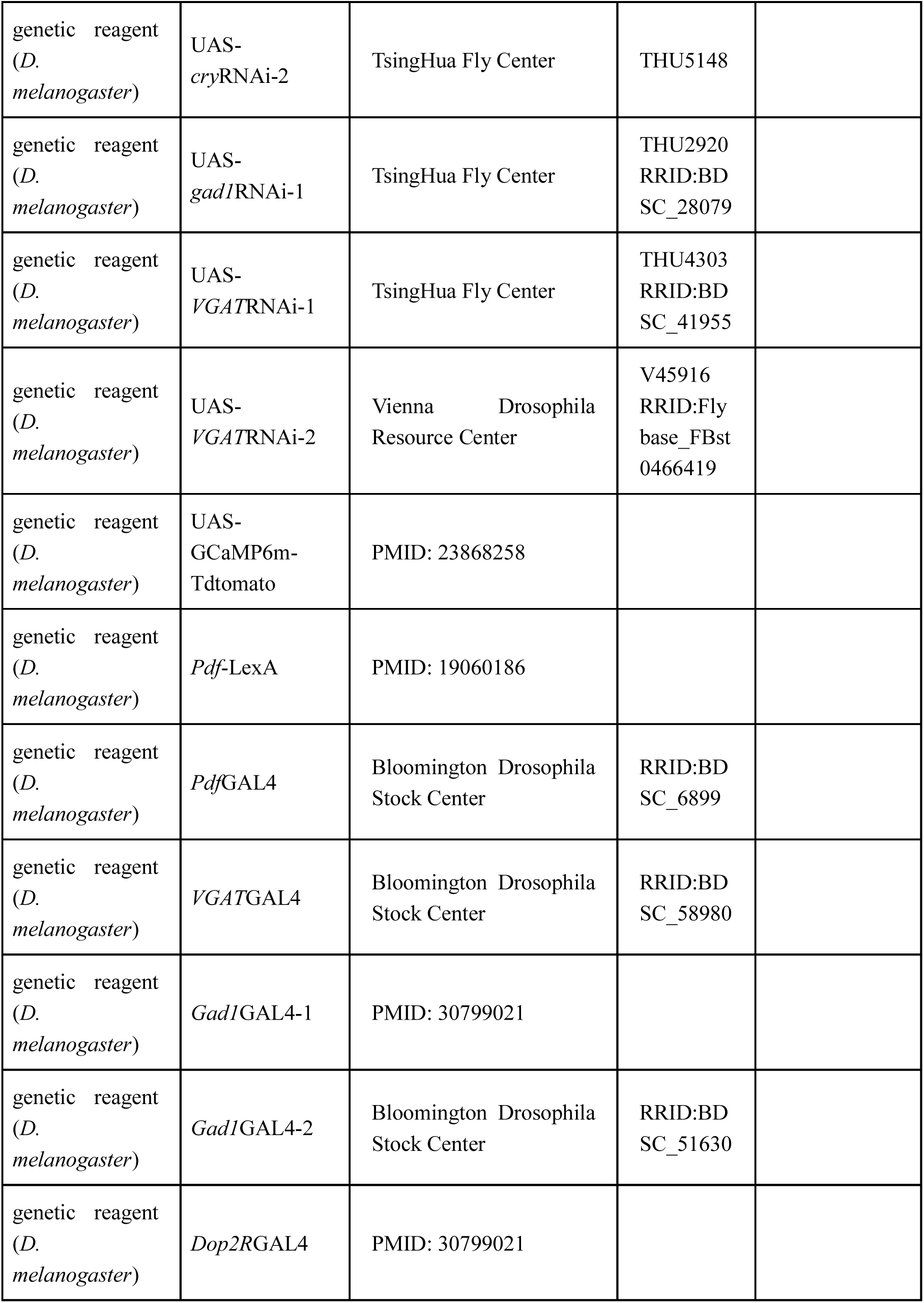
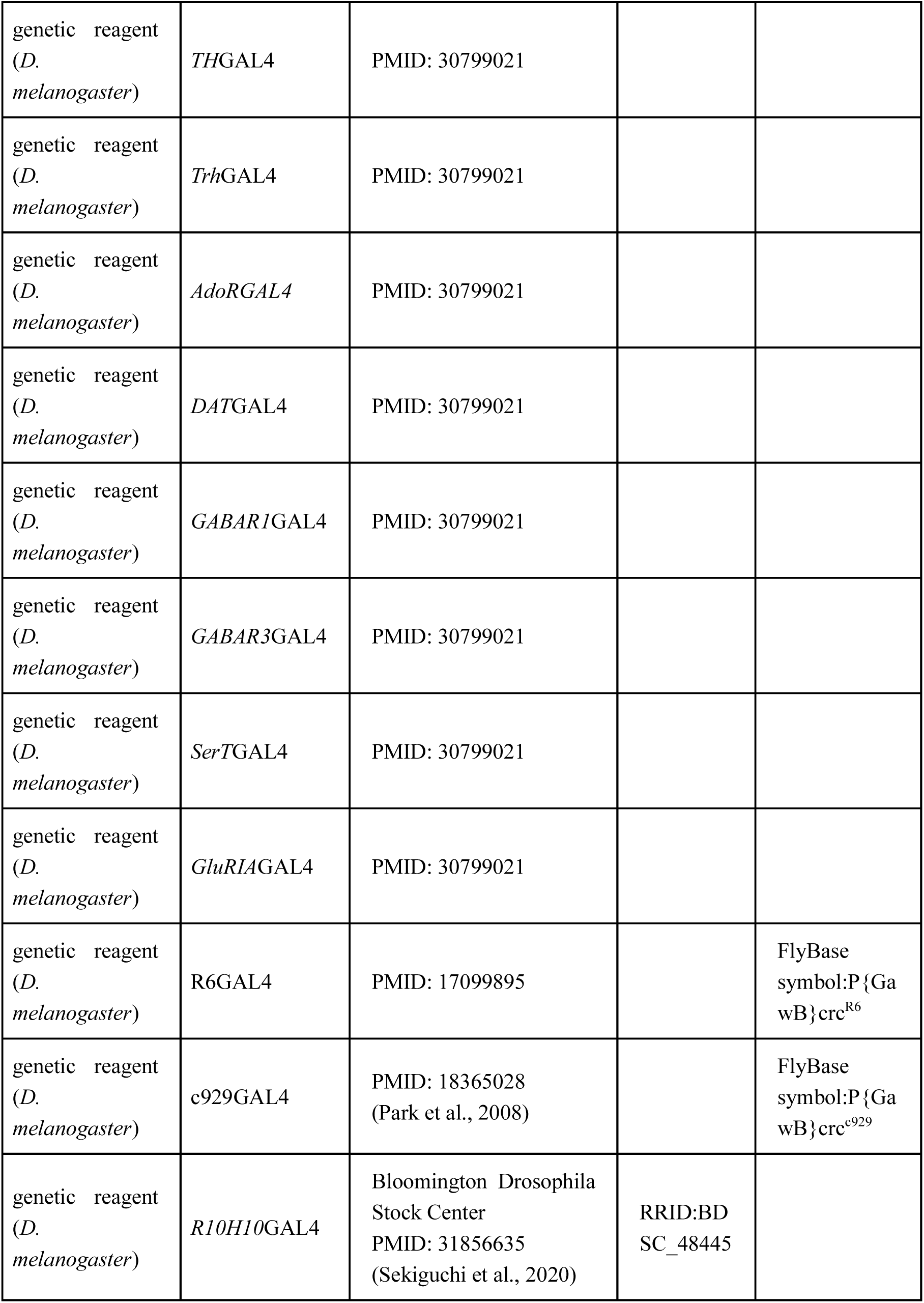
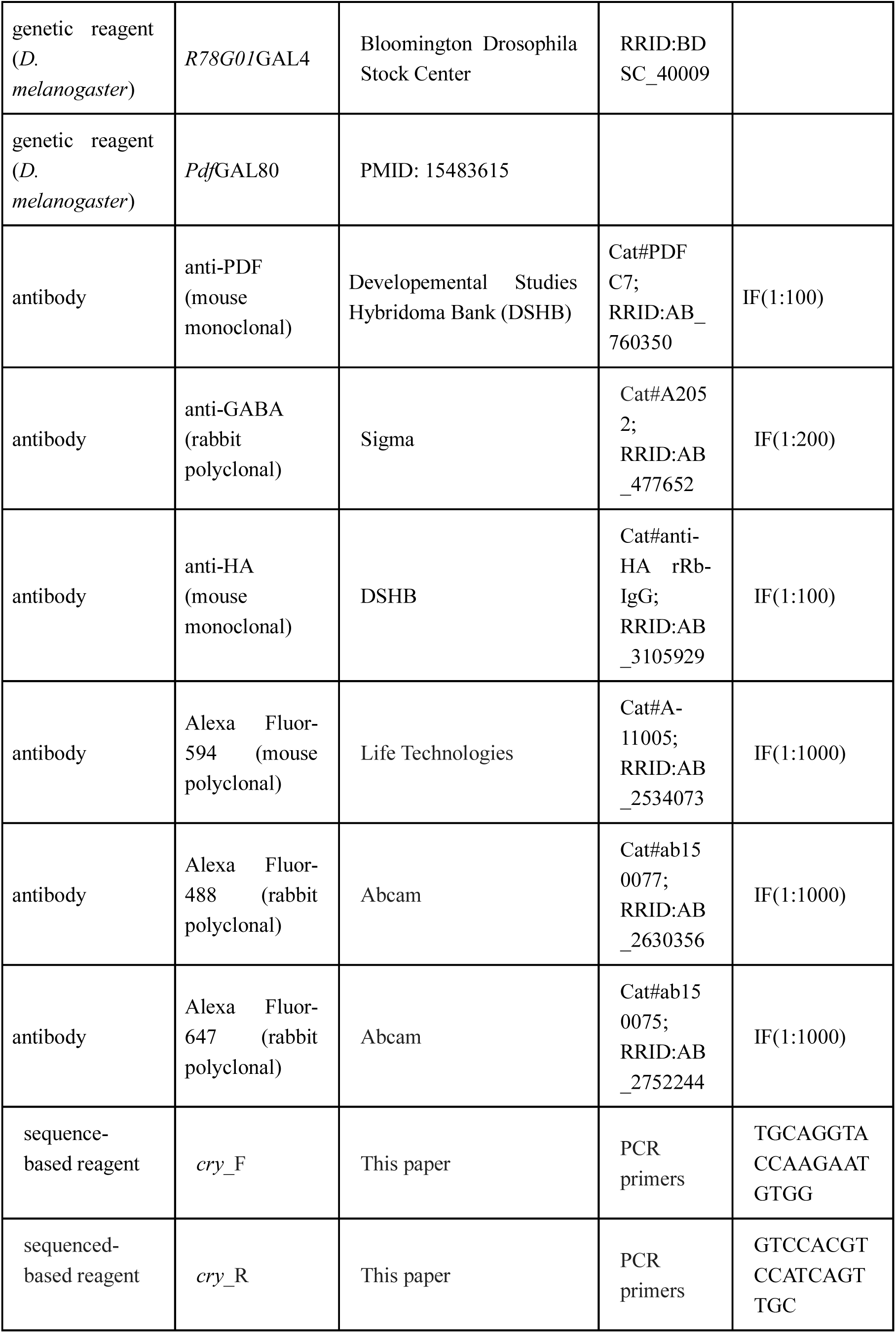
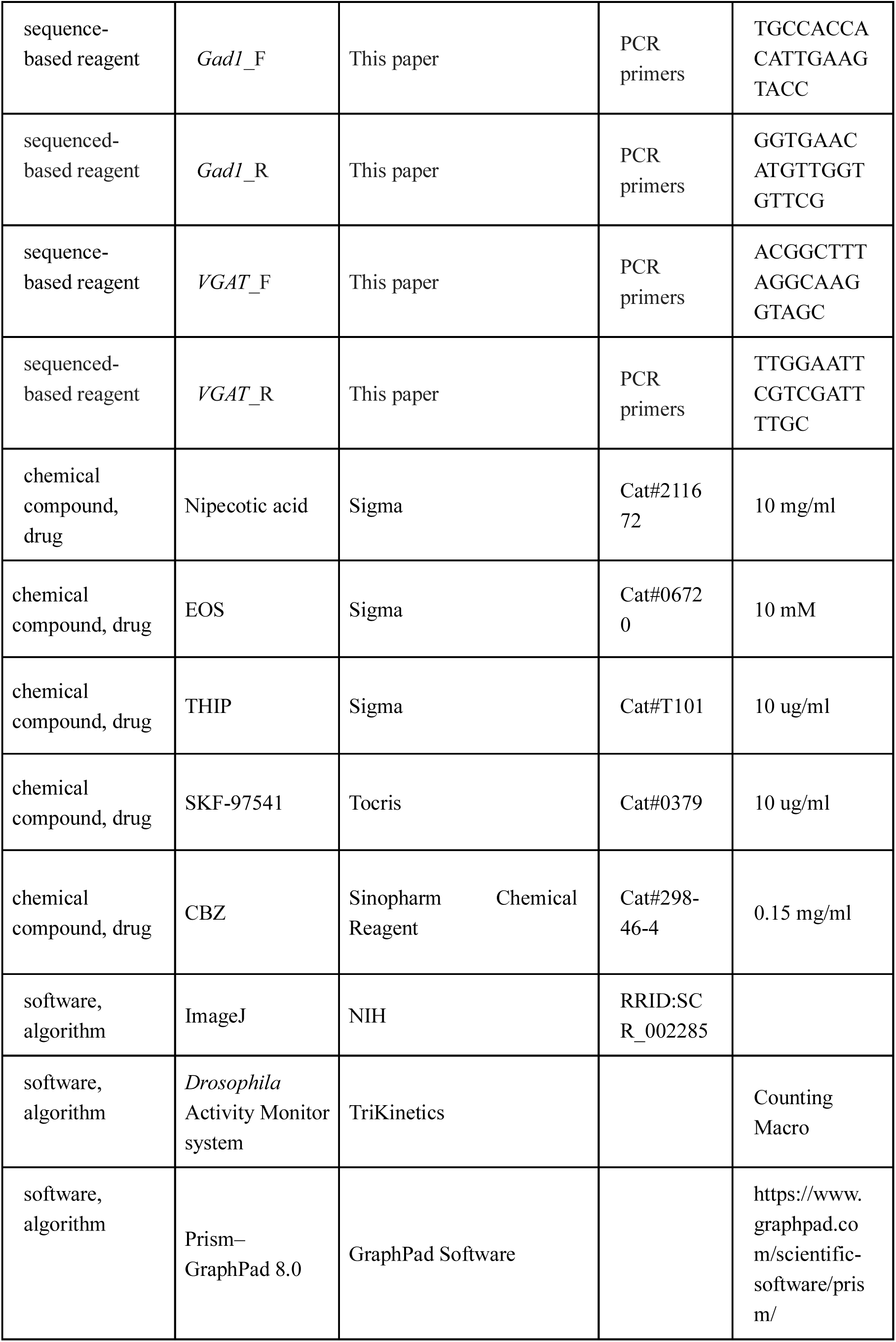
—key resources table.

### Figure Legend

**Figure 1—figure supplement 1.**
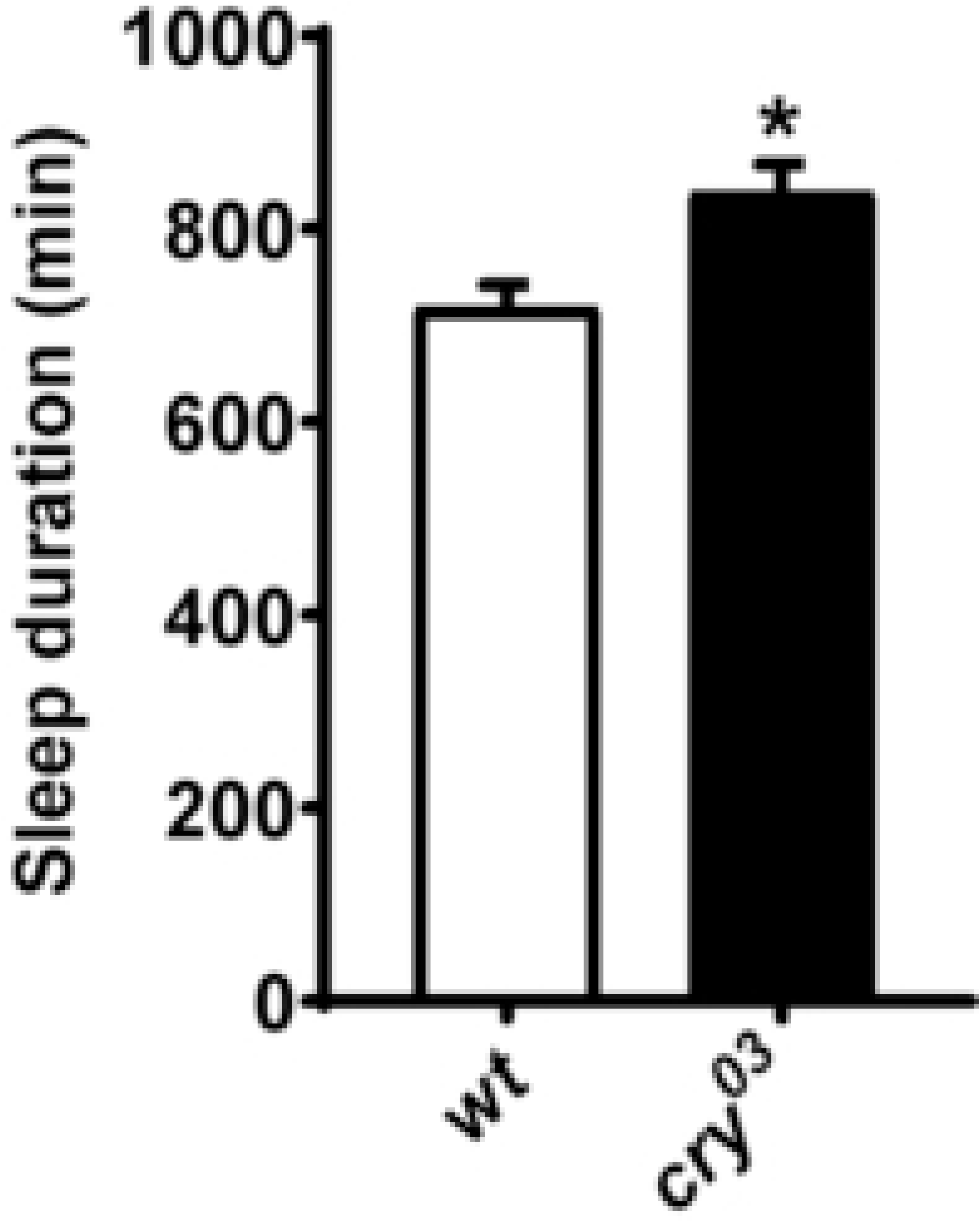
*cry* knock-out mutation lengthens sleep duration under 4L20D. Daily sleep duration of male WT flies and *cry* knock-out flies, monitored under 4L20D (n = 53, 35, 47). Student’s *t*-test: \**P* < 0.05. Error bars represent SEM.

**Figure 2—figure supplement 1.**
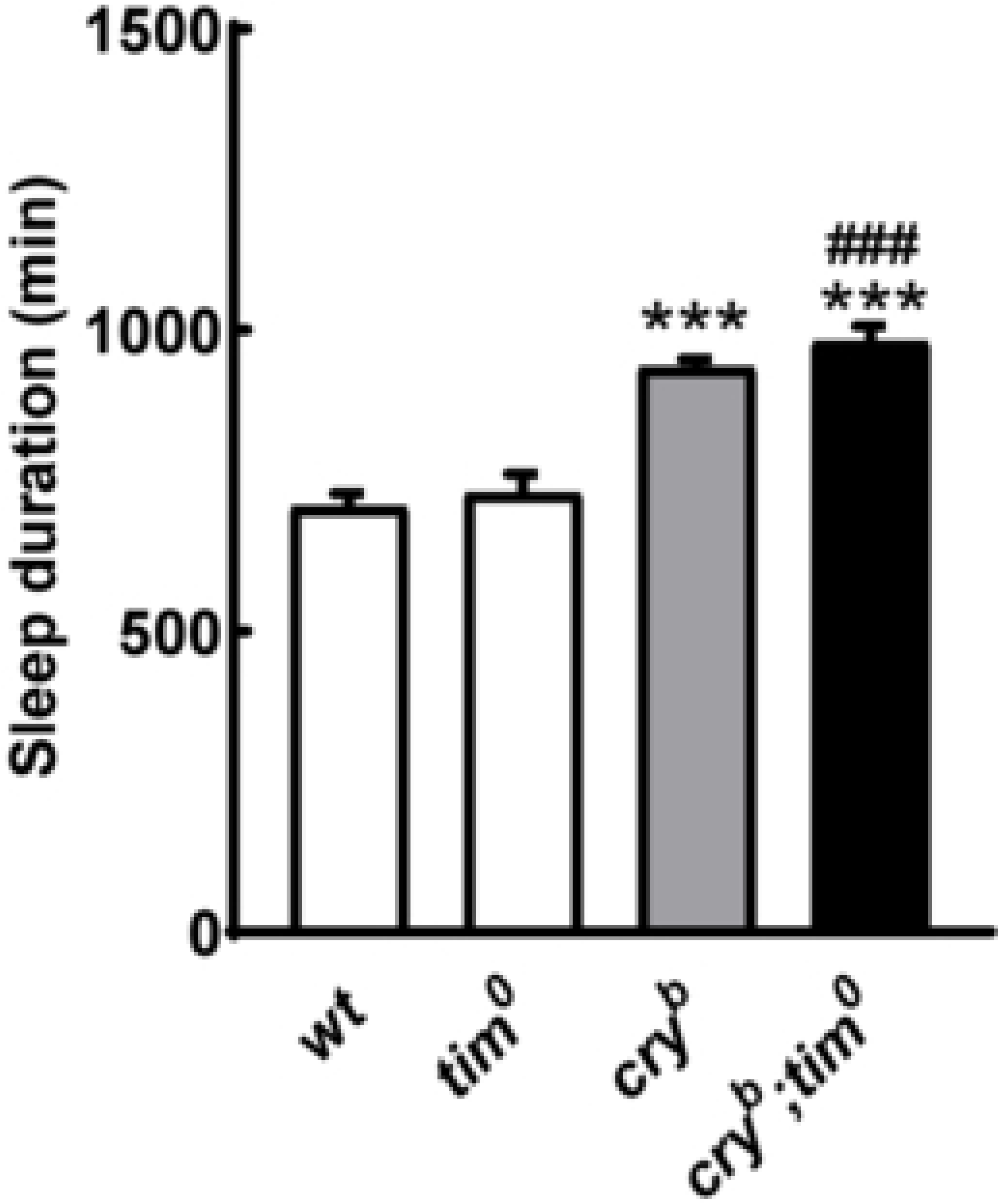
CRY does not regulate sleep duration via TIM. Daily sleep duration of male WT flies and flies mutant for *cry* and/or *tim*, monitored under 4L20D (n = 58, 63, 91, 33 flies). Two-tailed Student’s *t*-test: compared to WT background, \*\*\**P* < 0.001; compared to *tim^0^*, ###*P* < 0.001. Error bars represent SEM.

**Figure 2—figure supplement 2.**
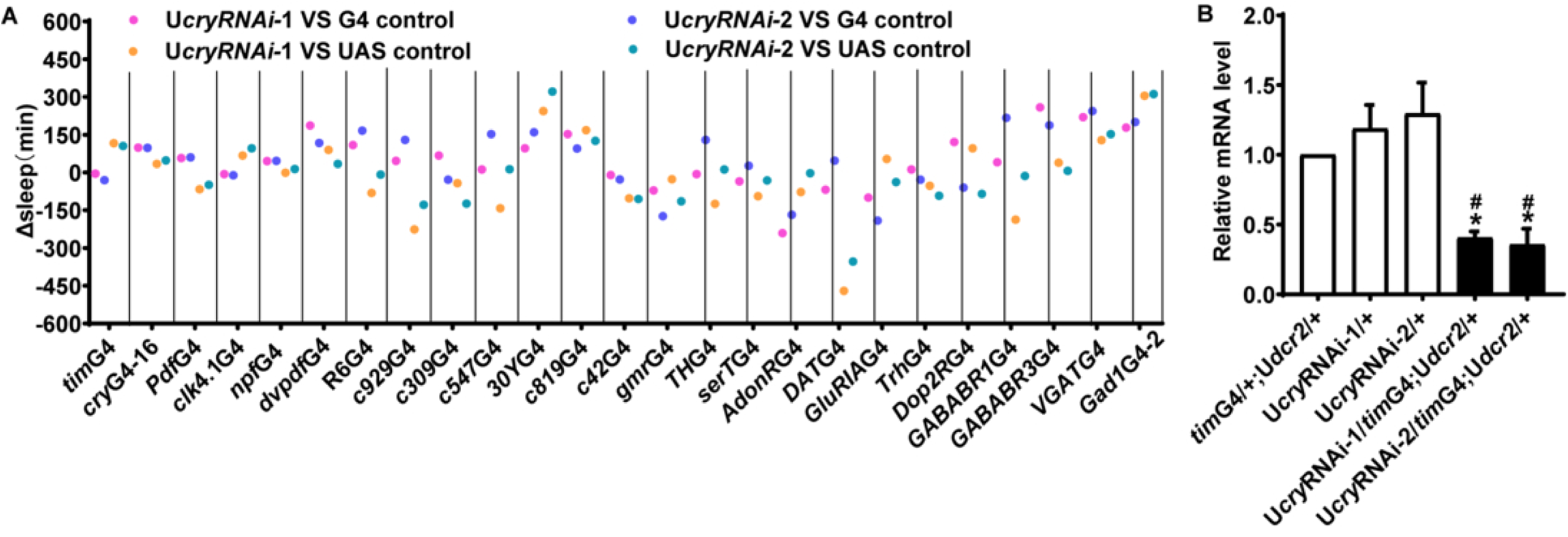
Screening for anatomical substrates that mediate the effects of CRY on sleep/wakefulness. (A) Difference in sleep duration between male *cry* RNAi flies compared to GAL4 and UAS controls under 4L20D (n = 20-92). (B) Plots of relative mRNA abundance for cry determined by qRT-PCR in whole-head extracts of *cry*RNAi and control flies under 4L20D (n = 4). The value of the GAL4 control group was set to 1. Mann-Whitney test: compared to GAL4 control, \**P* < 0.05; compared to UAS control, #*P* < 0.05. G4, GAL4; U, UAS.

**Figure 2—figure supplement 3.**
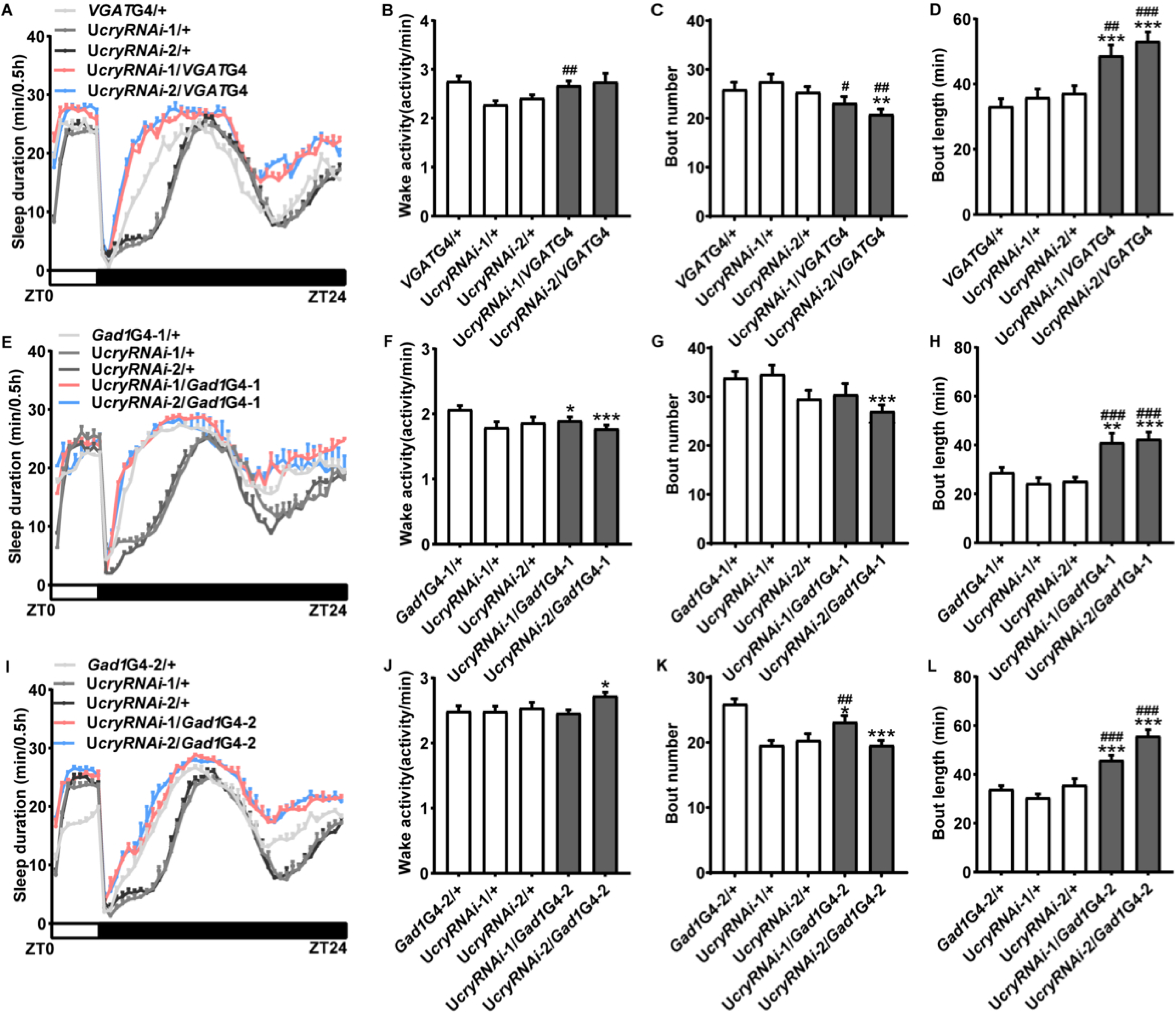
CRY promotes wakefulness in GABAergic neurons. (A-D) Sleep profile (A), daily waking activity (B), sleep bout number (C), average sleep bout length (D) of male flies with *cry* knocked down in GABAergic neurons by *VGAT*GAL4 monitored under 4L20D in Fig 2A. (E-H) Sleep profile (E), daily waking activity (F), sleep bout number (G), average sleep bout length (H) of male flies with *cry* knocked down in GABAergic neurons by *Gad1*GAL4-1 monitored under 4L20D in Fig 2B. (I-L) Sleep profile (I), daily waking activity (J), sleep bout number (K), average sleep bout length (L) of male flies with *cry* knocked down in GABAergic neurons by *Gad1*GAL4-2 monitored under 4L20D in Fig 2C.. One-way ANOVA with Bonferroni multiple comparison test: compared to GAL4 control, \*\*\**P* < 0.001; compared to UAS control, #*P* < 0.05, ###*P* < 0.001.

**Figure 2—figure supplement 4.**
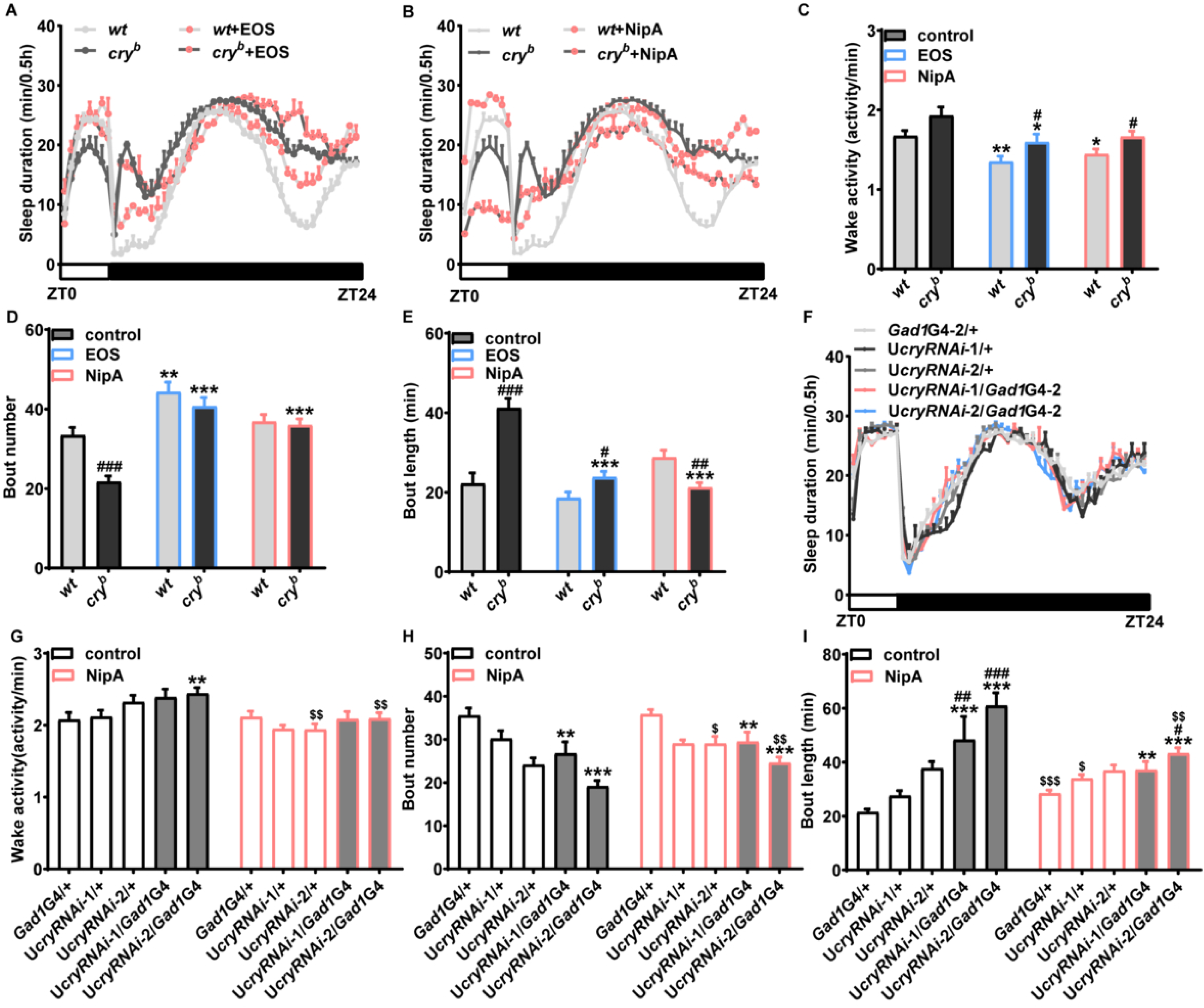
CRY promotes wakefulness via GABA signaling. (A, B) Sleep profile of male WT and *cry* mutant flies fed with NipA (A) or EOS (B) under 4L20D in Fig 2E. White box indicates light period while black box indicates dark period. (C-E) Daily waking activity (C), sleep bout number (D), average sleep bout length (E) of male WT and *cry* mutant flies fed with NipA or EOS under 4L20D in Fig 2E. Two-tailed Student’s *t*-test: compared to WT, #*P* < 0.05, ##*P* < 0.01, ###*P* < 0.001; compared to vehicle control, \**P* < 0.05, \*\**P* < 0.01, \*\*\**P* < 0.001. (F-I) Sleep profile (F), daily waking activity (G), sleep bout number (H), average sleep bout length (I) of *cry* RNAi and control flies fed with NipA under 4L20D in Fig 2F. One-way ANOVA with Bonferroni multiple comparison test: compared to GAL4 control, \*\**P* < 0.01, \*\*\**P* < 0.001; compared to UAS control, #*P* < 0.05, ##*P* < 0.01, ###*P* < 0.001. For comparing to vehicle control, two-tailed Student’s *t*-test was used, $*P* < 0.05, $$*P* < 0.01, $$$*P* < 0.001. Error bars represent SEM. G4, GAL4; U, UAS.

**Figure 2—figure supplement 5.**
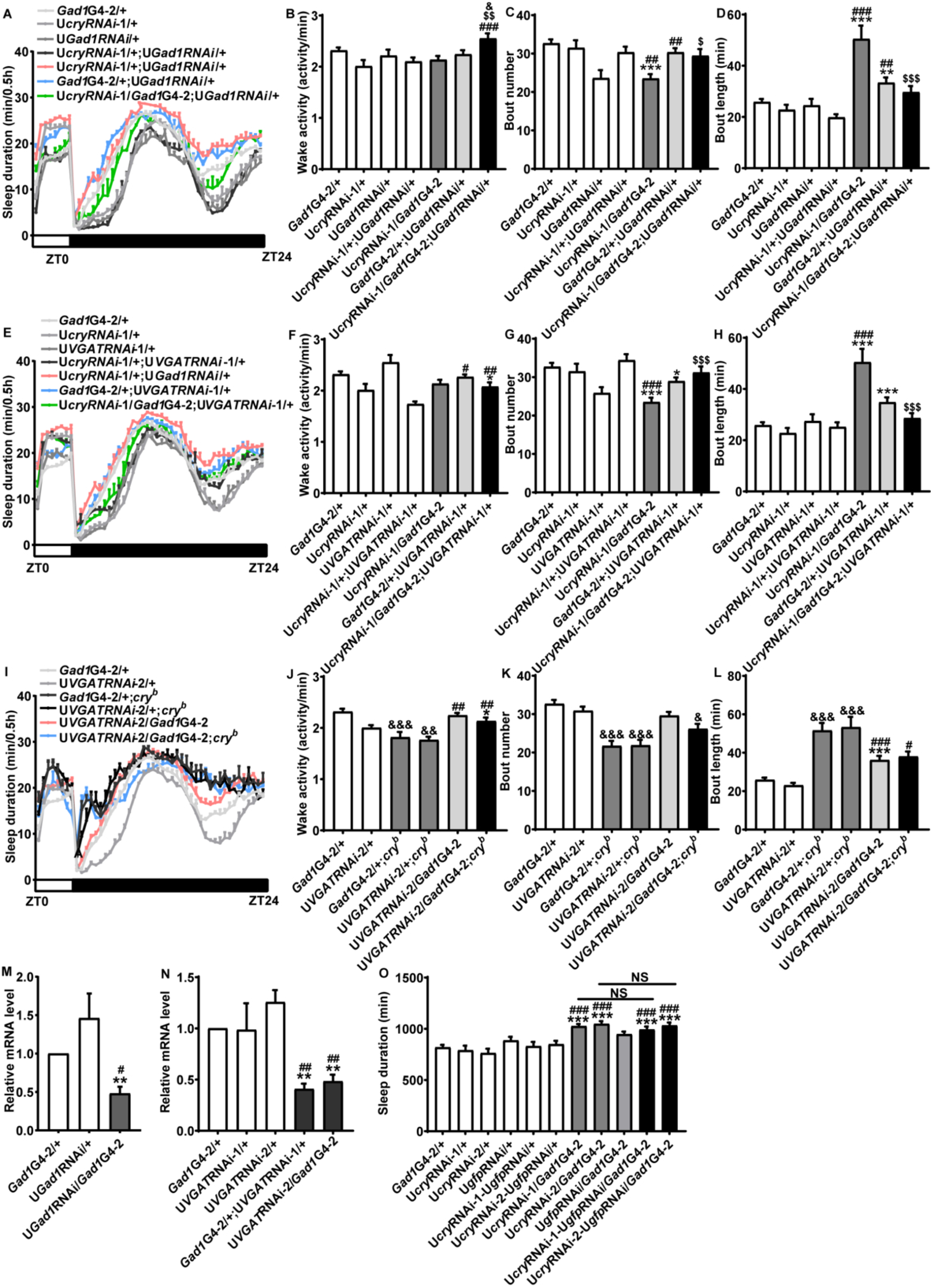
CRY promotes wakefulness via GABA-related genes. (A-D) Sleep profile (A), daily waking activity (B), sleep bout number (C), average sleep bout length (D) of male flies with *cry* and *gad1* knocked down in GABAergic neurons monitored under 4L20D in Fig 2G. (E-H) Sleep profile (E), daily waking activity (F), sleep bout number (G), average sleep bout length (H) of male flies with *cry* and *VGAT* knocked down in GABAergic neurons monitored under 4L20D in Fig 2H. For comparison between RNAi flies vs. UAS/GAL4 controls, one-way ANOVA with Bonferroni multiple comparison test was used: compared to GAL4 control, \**P* < 0.05, \*\**P* < 0.01, \*\*\**P* < 0.001; compared to UAS control, #*P* < 0.05, ##*P* < 0.01, ###*P* < 0.001. For comparing to RNAi control, two-tailed Student’s *t*-test was used: compared to UAS*cry*RNAi/GAL4 control, $*P* < 0.05, $$*P* < 0.01, $$$*P* < 0.001; compared to UAS*Gad1*RNAi/GAL4 or U*VGAT*RNAi/GAL4 control, &&&*P* < 0.001. (I-L) Sleep profile (I), daily waking activity (J), sleep bout number (K), average sleep bout length (L) of *cry* mutant flies with *VGAT* knocked down in GABAergic neurons monitored under 4L20D in Fig 2I. For comparison between RNAi flies vs. UAS/GAL4 controls, one-way ANOVA with Bonferroni multiple comparison test was used: compared to GAL4 control, \**P* < 0.05, \*\*\**P* < 0.001; compared to UAS control, #*P* < 0.05, ##*P* < 0.01, ###*P* < 0.001. For comparison between mutant vs. control, two-tailed Student’s *t*-test was used: compared to WT background, &*P* < 0.05, &&*P* < 0.01, &&&*P* < 0.001. (M, N) Plots of relative mRNA abundance for *Gad1* (M) and *VGAT* (N) determined by qRT-PCR in whole-head extracts of RNAi and control flies under 4L20D (n = 6). The value of the GAL4 control group was set to 1. Mann-Whitney test: compared to GAL4 control, \*\**P* < 0.01; compared to UAS control, ##*P* < 0.01. (O) Daily sleep duration of male flies with *cry* and *gfp* knocked down in GABAergic neurons by *Gad1*GAL4 monitored under 4L20D (n = 72, 26, 29, 20, 30, 31, 68, 32, 31, 29, 29). One-way ANOVA with Bonferroni multiple comparison test: compared to GAL4 control, \*\*\**P* < 0.001; compared to UAS control, #*P* < 0.05, ###*P* < 0.001. Error bars represent SEM; G4, GAL4; U, UAS; NS, not significant.

**Figure 3—figure supplement 1.**
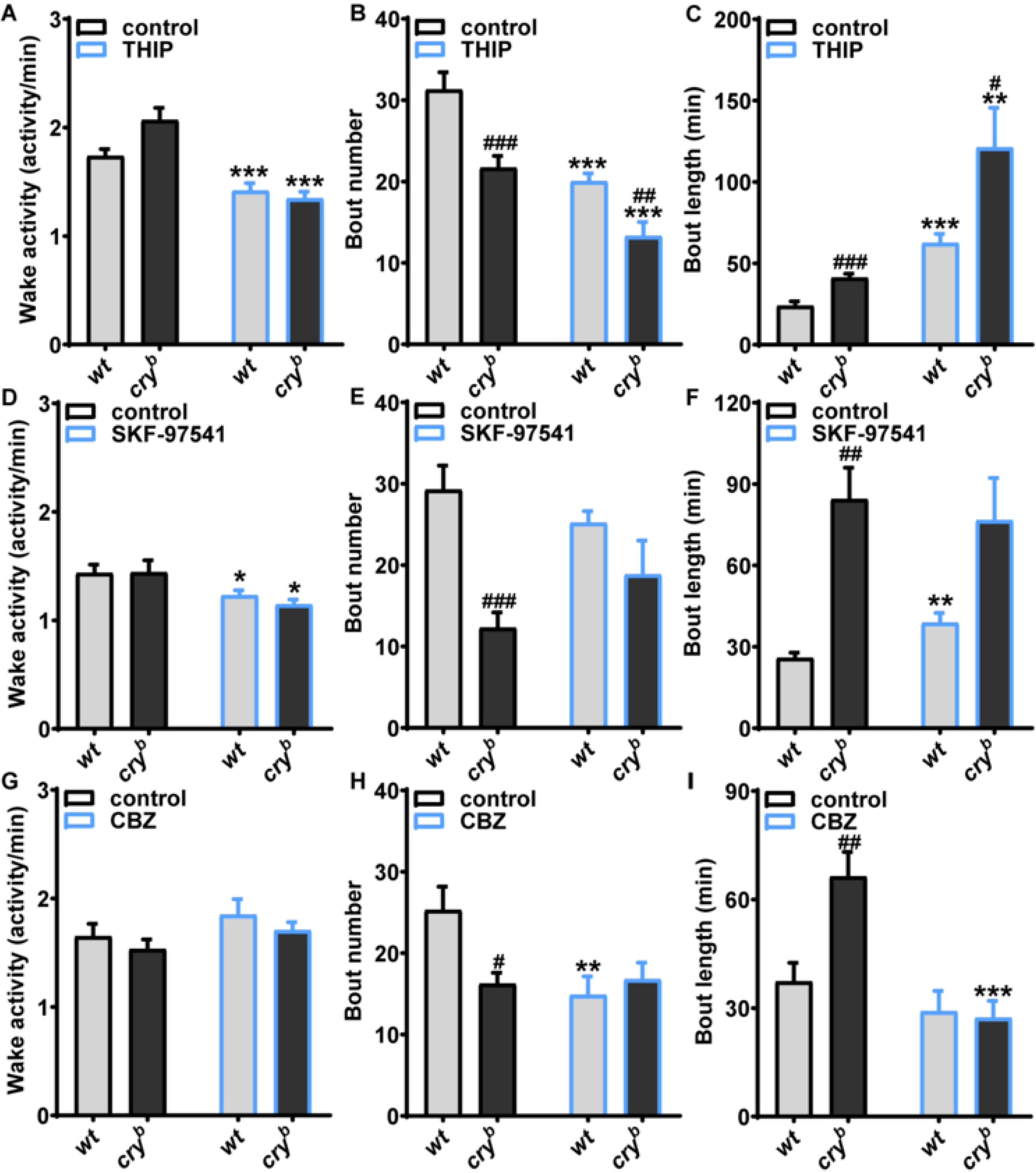
GABA-A receptor mediates the effects of *cry* mutation on sleep/wakefulness. (A-C) Daily waking activity (A), sleep bout number (B), average sleep bout length (C) of male WT and *cry* mutant flies fed with THIP under 4L20D in Fig 3A. (D-F) Daily waking activity (D), sleep bout number (E), average sleep bout length (F) of male WT and *cry* mutant flies fed with SKF-97541 under 4L20D in Fig 3C. (G-I) Daily waking activity (G), sleep bout number (H), average sleep bout length (I) of male WT and *cry* mutant flies fed with CBZ under 4L20D in Fig 3D. Two-tailed Student’s *t*-test: compared to WT, #*P* < 0.05, ##*P* < 0.01, ###*P* < 0.001; compared to vehicle control, \**P* < 0.05, \*\**P* < 0.01, \*\*\**P* < 0.001. Error bars represent SEM.

**Figure 3—figure supplement 2.**
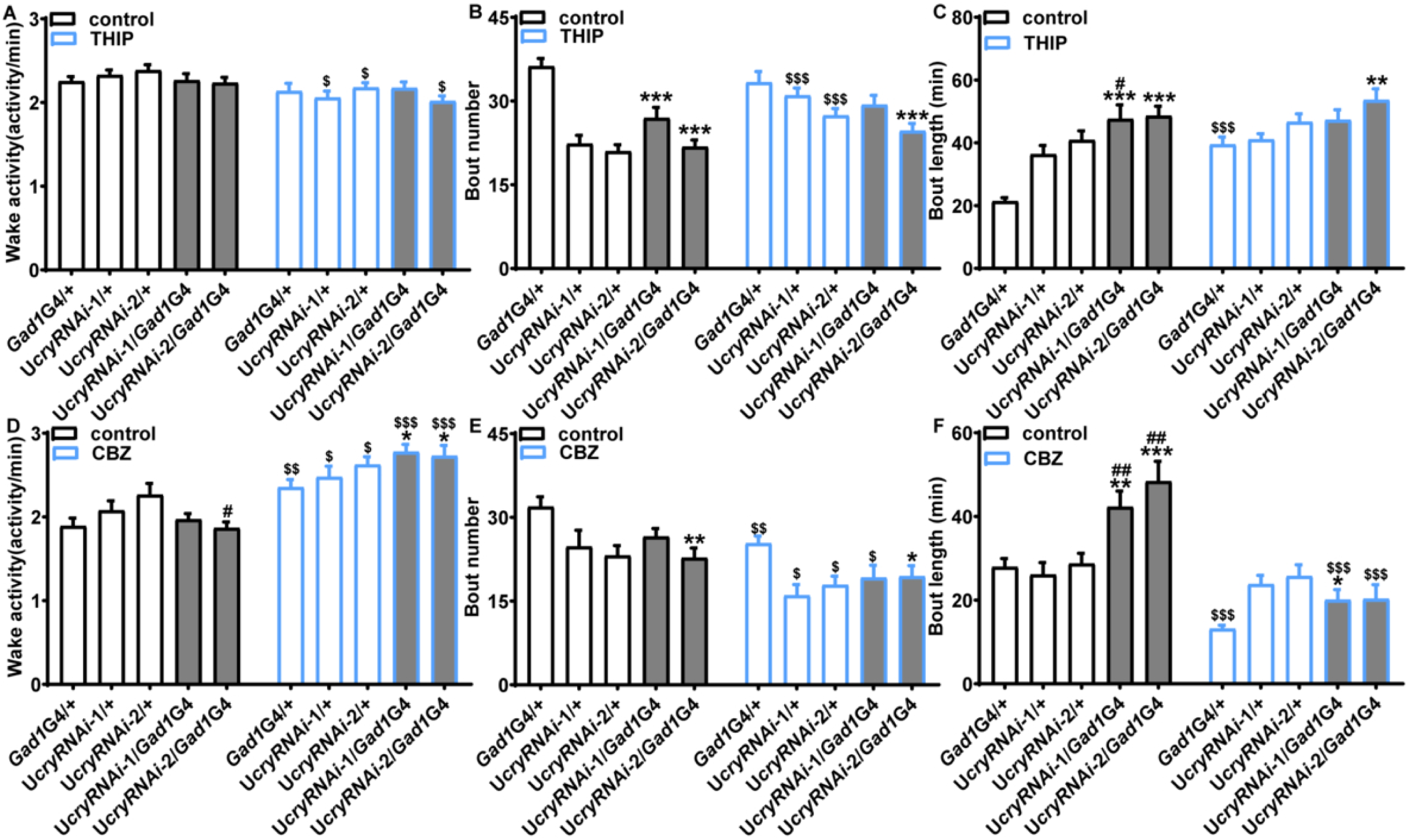
GABA-A receptor mediates the effects of *cry* RNAi flies on sleep/wakefulness. (A-C) Daily waking activity (A), sleep bout number (B), average sleep bout length (C) of male *cry* RNAi and control flies fed with THIP under 4L20D in Fig 3F. (D-F) Daily waking activity (D), sleep bout number (E), average sleep bout length (F) of male *cry* RNAi and control flies fed with CBZ under 4L20D in Fig 3H. For comparison between RNAi flies vs. UAS/GAL4 controls, one-way ANOVA with Bonferroni multiple comparison test was used: compared to GAL4 control, \**P* < 0.05, \*\**P* < 0.01, \*\*\**P* < 0.001; compared to UAS control, #*P* < 0.05, ##*P* < 0.01. For comparing to vehicle control, two-tailed Student’s *t*-test was used, $*P* < 0.05, $$*P* < 0.01, $$$*P* < 0.001. Error bars represent SEM. G4, GAL4; U, UAS.

**Figure 3—figure supplement 3.**
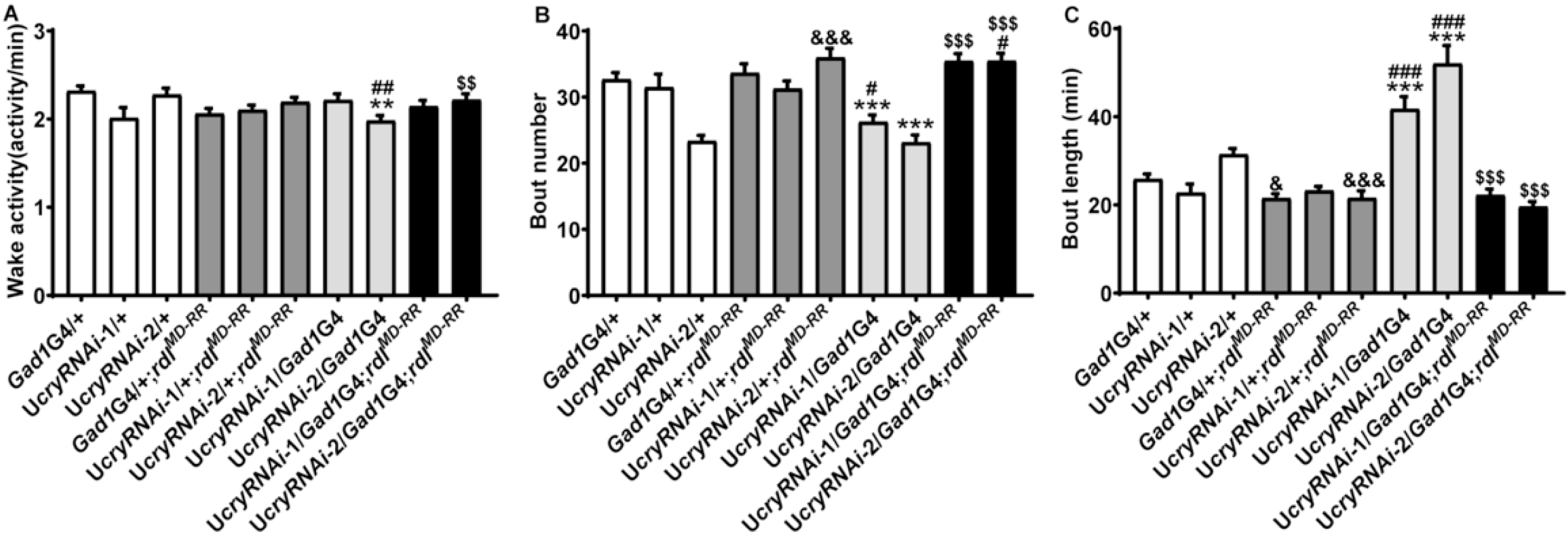
GABA-A receptor mediates the effects of *cry* RNAi flies on sleep/wakefulness. (A-C) Daily waking activity (A), sleep bout number (B), average sleep bout length (C) of male *rdl^MD-RR^*flies with *cry* knocked down in GABAergic neurons under 4L20D in Fig 3J. For comparison between RNAi flies vs. UAS/GAL4 controls, one-way ANOVA with Bonferroni multiple comparison test was used: compared to GAL4 control, \*\**P* < 0.01, \*\*\**P* < 0.001; compared to UAS control, #*P* < 0.05, ##*P* < 0.01, ###*P* < 0.001. For comparing to WT or UAS*cry*RNAi/GAL4 control, two-tailed Student’s *t*-test was used: compared to WT, &*P* < 0.05; &&&*P* < 0.001; compared to UAS*cry*RNAi/GAL4 control, $$*P* < 0.01, $$$*P* < 0.001. Error bars represent SEM. G4, GAL4; U, UAS.

**Figure 4—figure supplement 1.**
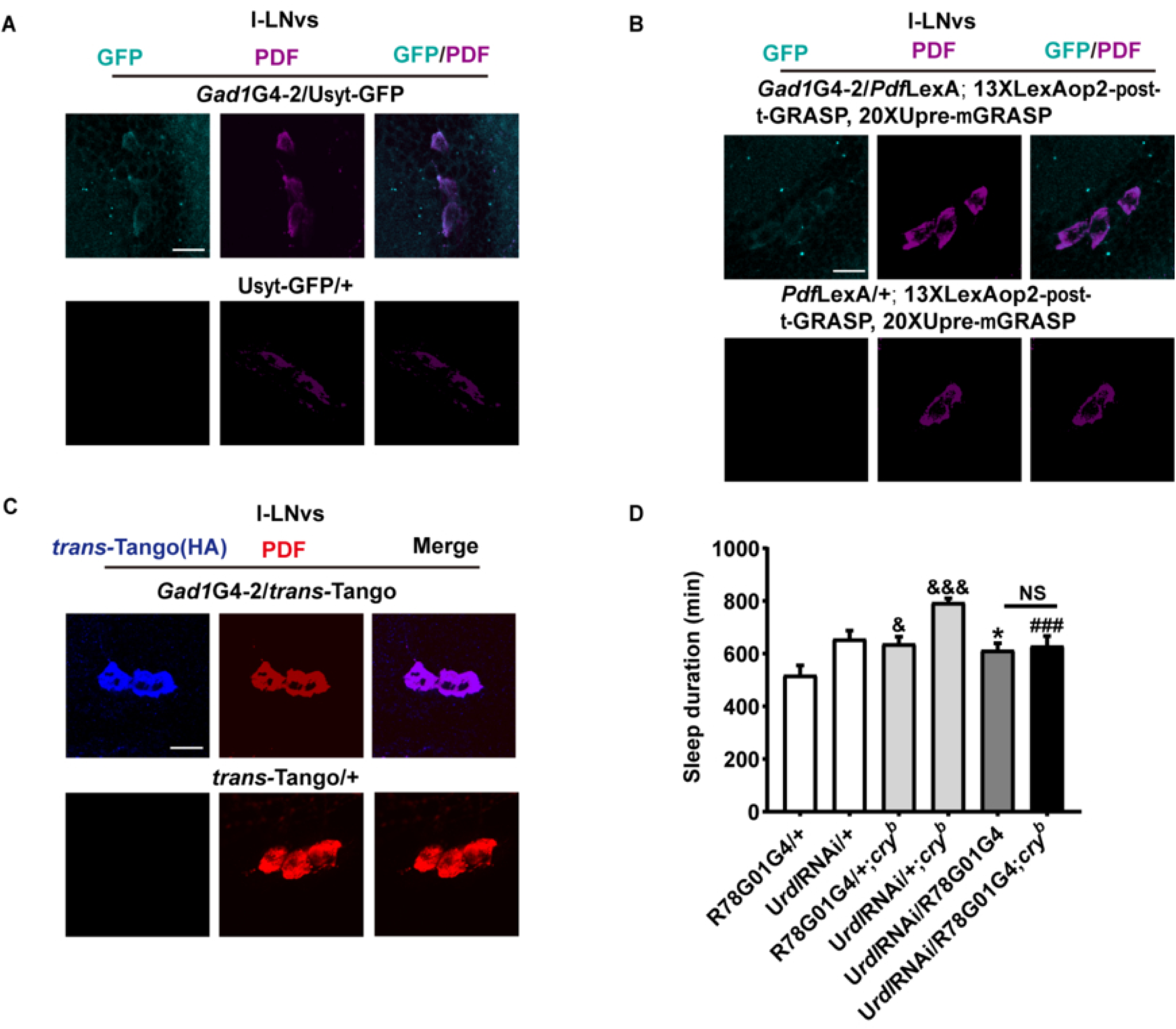
GABAergic neurons project to and form synapses with the l-LNvs. (A) l-LNvs of flies expressing syt-GFP in GABAergic neurons using *Gad1*Gal4 maintained under 4L20D and immunostained with PDF antisera. (B) l-LNvs of flies expressing Pre-mGRASP in GABAergic neurons using *Gad1*Gal4 and t-GRASP in the l-LNvs using *Pdf*LexA maintained under 4L20D. Brains are immunostained with PDF antisera. (C) l-LNvs of flies expressing trans-Tango in GABAergic neurons using *Gad1*Gal4 maintained under 4L20D and immunostained with HA and PDF antisera. (D) Daily sleep duration of *cry* mutant flies with *rdl* knocked down in the l-LNvs using R78G01GAL4 and relevant controls, monitored under 4L20D (n = 31, 31, 64, 32, 31, 32 flies). For comparison between RNAi flies vs. UAS/GAL4 controls, one-way ANOVA with Bonferroni multiple comparison test was used: compared to GAL4 control, \**P* < 0.05; compared to UAS control, ###*P* < 0.001. For comparison between mutant vs. control, two-tailed Student’s *t*-test was used: compared to WT background, &*P* < 0.05, &&&*P* < 0.001; compared to U*rdl*RNAi/GAL4, not significant. The scale bar represents 15 µm. Error bars represent SEM. G4, GAL4; U, UAS; LexAop, LexA operator.

**Figure 4—figure supplement 2.**
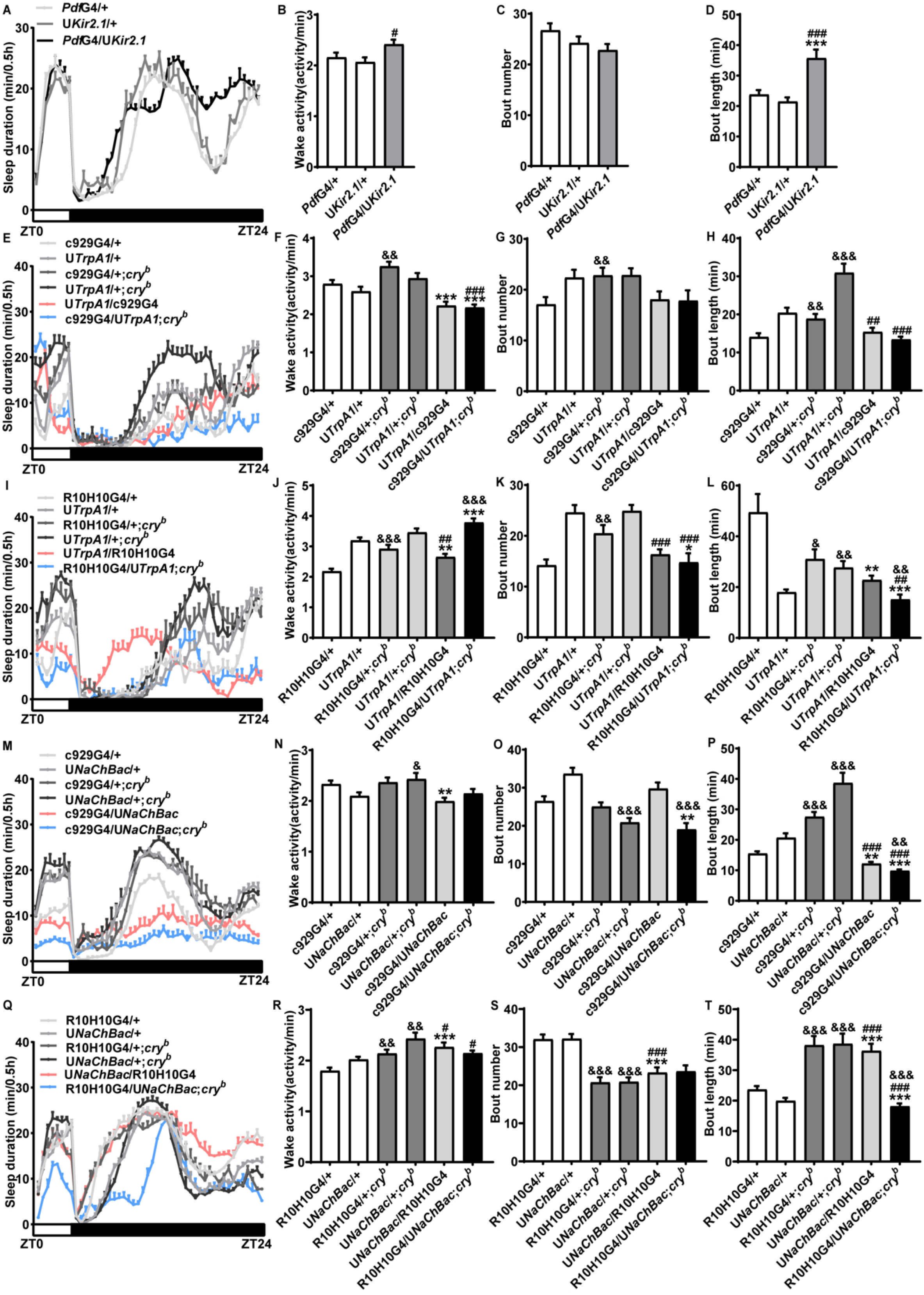
CRY promote wakefulness by impinging on the excitability of l-LNv. (A-D) Sleep profile (A), daily waking activity (B), sleep bout number (C), average sleep bout length (D) of male flies expressing Kir2.1 in PDF neuron using *Pdf*GAL4 and controls, monitored under 4L20D in Fig 4G. White box indicates light period while black box indicates dark period. (E-H) Sleep profile (E), daily waking activity (F), sleep bout number (G), average sleep bout length (H) of male *cry* mutant flies expressing TrpA1 in the l-LNvs using c929GAL4 and relevant controls in Fig 4H and I, monitored under 4L20D and 29℃ to activate TrpA1. (I-L) Sleep profile (I), daily waking activity (J), sleep bout number (K), average sleep bout length (L) of male *cry* mutant flies expressing TrpA1 in the l-LNvs using R10H10GAL4 and relevant controls in Fig 4I, monitored under 4L20D and 29℃ to activate TrpA1. For comparison with UAS/GAL4 controls, one-way ANOVA with Bonferroni multiple comparison test was used: compared to GAL4 control, \**P* < 0.05, \*\**P* < 0.01, \*\*\**P* < 0.001; compared to UAS control, #*P* < 0.05, ##*P* < 0.01, ###*P* < 0.001. For comparison between mutant vs. control, two-tailed Student’s *t*-test was used: &*P* < 0.05, &&*P* < 0.01, &&&*P* < 0.001. (M-P) Sleep profile (M), daily waking activity (N), sleep bout number (O), average sleep bout length (P) of *cry* mutant flies expressing NachBac in the l-LNvs using c929GAL4 and relevant controls in Fig 4J, monitored under 4L20D. (Q-T) Sleep profile (Q), daily waking activity (R), sleep bout number (S), average sleep bout length (T) of *cry* mutant flies expressing NachBac in the l-LNvs using R10H10GAL4 and relevant controls in Fig 4K, monitored under 4L20D. For comparison with UAS/GAL4 controls, one-way ANOVA with Bonferroni multiple comparison test was used: compared to GAL4 control, \*\**P* < 0.01, \*\*\**P* < 0.001; compared to UAS control, #*P* < 0.05, ###*P* < 0.001. For comparison between mutant vs. control, two-tailed Student’s *t*-test was used: &*P* < 0.05, &&*P* < 0.01, &&&*P* < 0.001. Error bars represent SEM. G4, GAL4; U, UAS.

**Figure 4—figure supplement 3.**
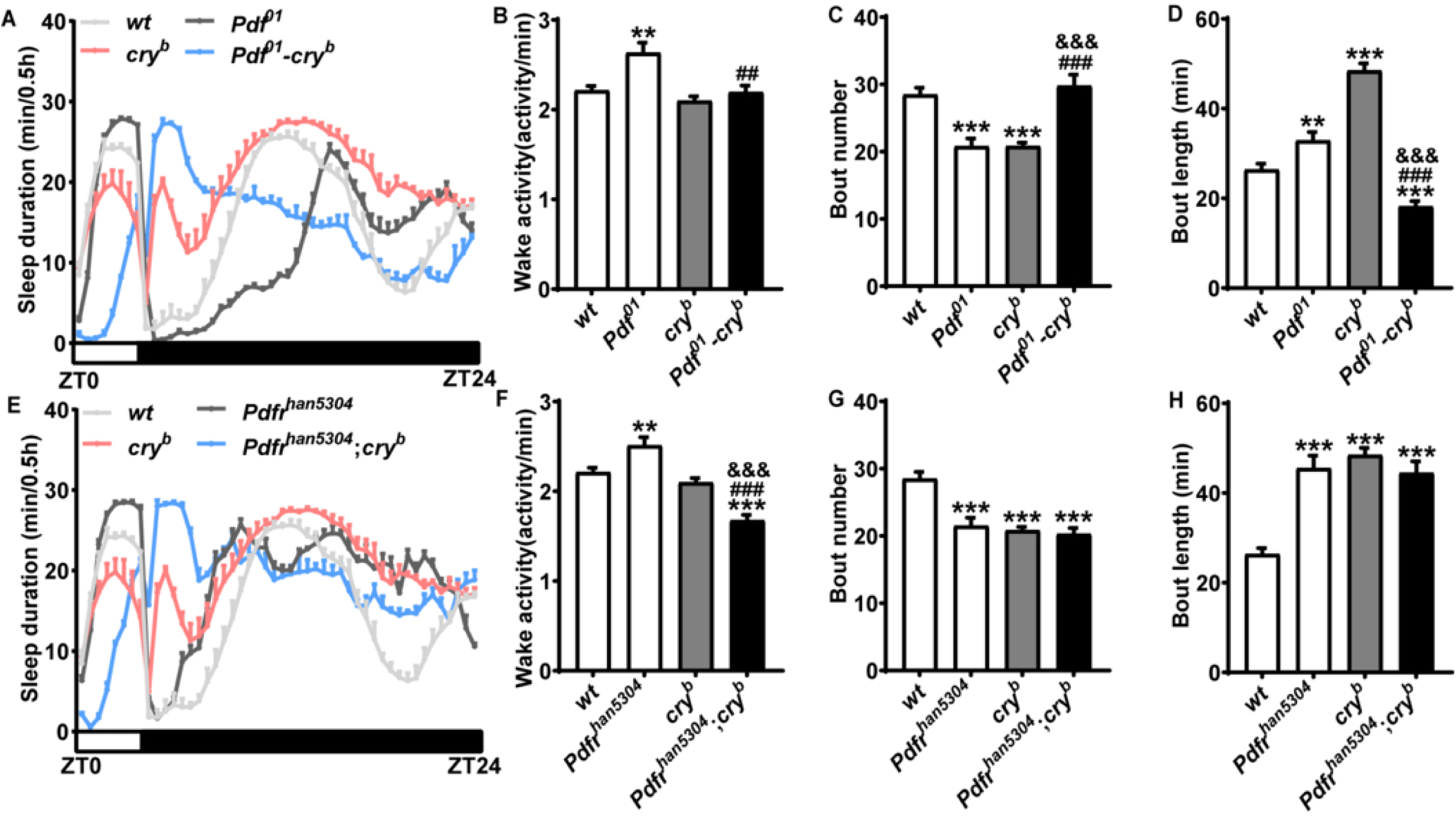
CRY promote wakefulness by impinging on PDF signaling. (A-D) sleep profile (A), daily waking activity (B), sleep bout number (C), average sleep bout length (D) of male *Pdf^01^-cry^b^* mutants along with relevant controls in Fig 4L, monitored under 4L20D. (E-H) Sleep profile (E), daily waking activity (F), sleep bout number (G), average sleep bout length (H) of male *Pdfr^han5304^*;*cry^b^*mutants along with relevant controls in Fig 4M, respectively, monitored under 4L20D. Two-tailed Student’s *t*-test: compared to WT background, \*\**P* < 0.01, \*\*\**P* < 0.001; compared to *Pdf^01^*or *Pdfr^han5304^*, ##*P* < 0.01, ###*P* < 0.001; compared to *cry^b^*, &&&*P* < 0.001. Error bars represent SEM.

**Figure 5—figure supplement 1.**
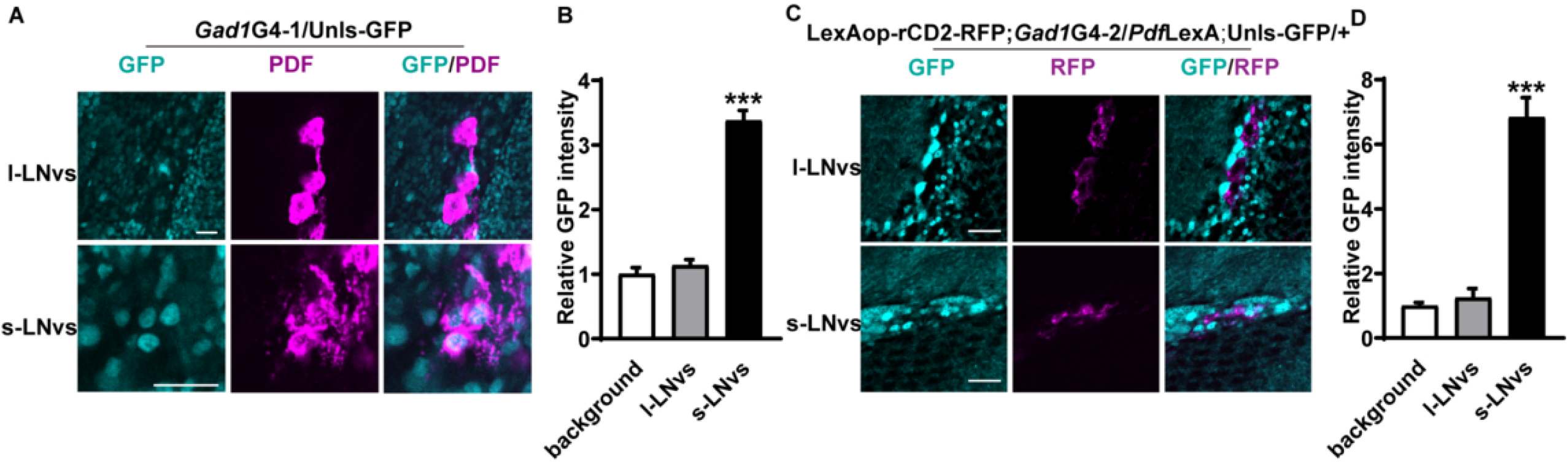
*Gad1*GAL4 drives expression in the s-LNvs but not l-LNvs. (A) Brains of male flies expressing nls-GFP driven by *Gad1*GAL4 maintained under 4L20D and immunostained with PDF antisera. (B) Quantification of GFP signal intensity of PDF neuron (n = 15, 18, 25 cells). Two-tailed Student’s *t* test: \*\*\**P* < 0.001. (C) Brains of male flies expressing nls-GFP in GABAergic neurons using *Gad1*Gal4 and rCD2-RFP in the PDF neurons using *Pdf*LexA maintained under 4L20D. (D) Quantification of GFP signal intensity of PDF neuron (n = 18, 22, 21 cells). Two-tailed Student’s *t* test: \*\*\**P* < 0.001. The scale bar represents 15 µm.

**Figure 5—figure supplement 2.**
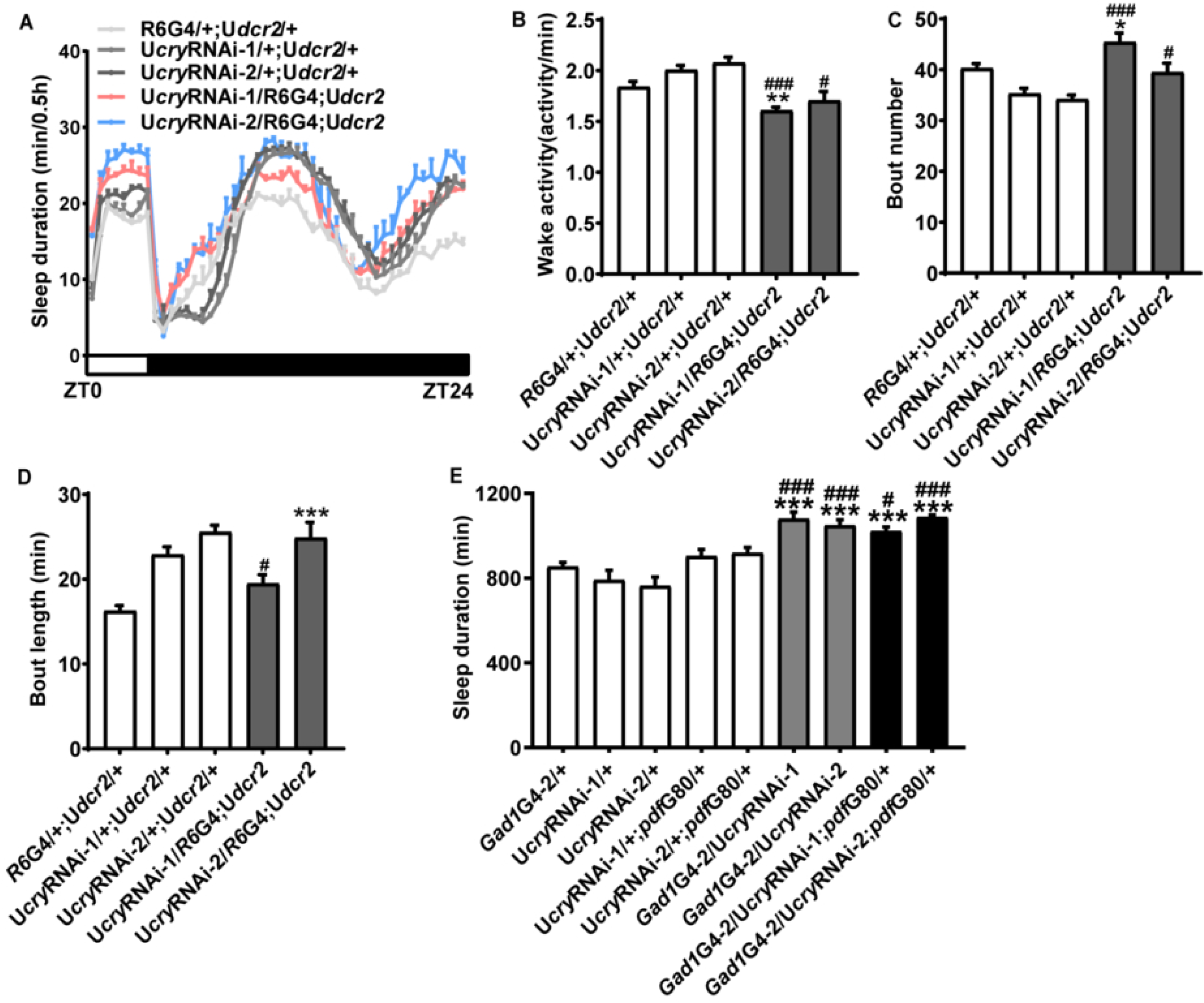
*cry* expression in the s-LNvs is necessary but not sufficient for maintaining normal sleep/wakefulness. (A-D) Sleep profile (A), daily waking activity (B), sleep bout number (C), average sleep bout length (D) of *cry* knocked down in s-LNvs by R6GAL4 monitored under 4L20D in Fig 5C. White box indicates light period while black box indicates dark period. (E) Daily sleep duration of *cry* knocked down in PDF-GABAergic neurons using *Gad1*GAL4 and *Pdf*GAL80, monitored under 4L20D (n =101, 26, 29, 27, 31, 23, 32, 73, 63). One-way ANOVA with Bonferroni multiple comparison test: compared to GAL4 control, \**P* < 0.05, \*\**P* < 0.01, \*\*\**P* < 0.001; compared to UAS control, #*P* < 0.05, ###*P* < 0.001. Error bars represent SEM. G4, GAL4; G80, GAL80; U, UAS.

**Figure 5—figure supplement 3.**
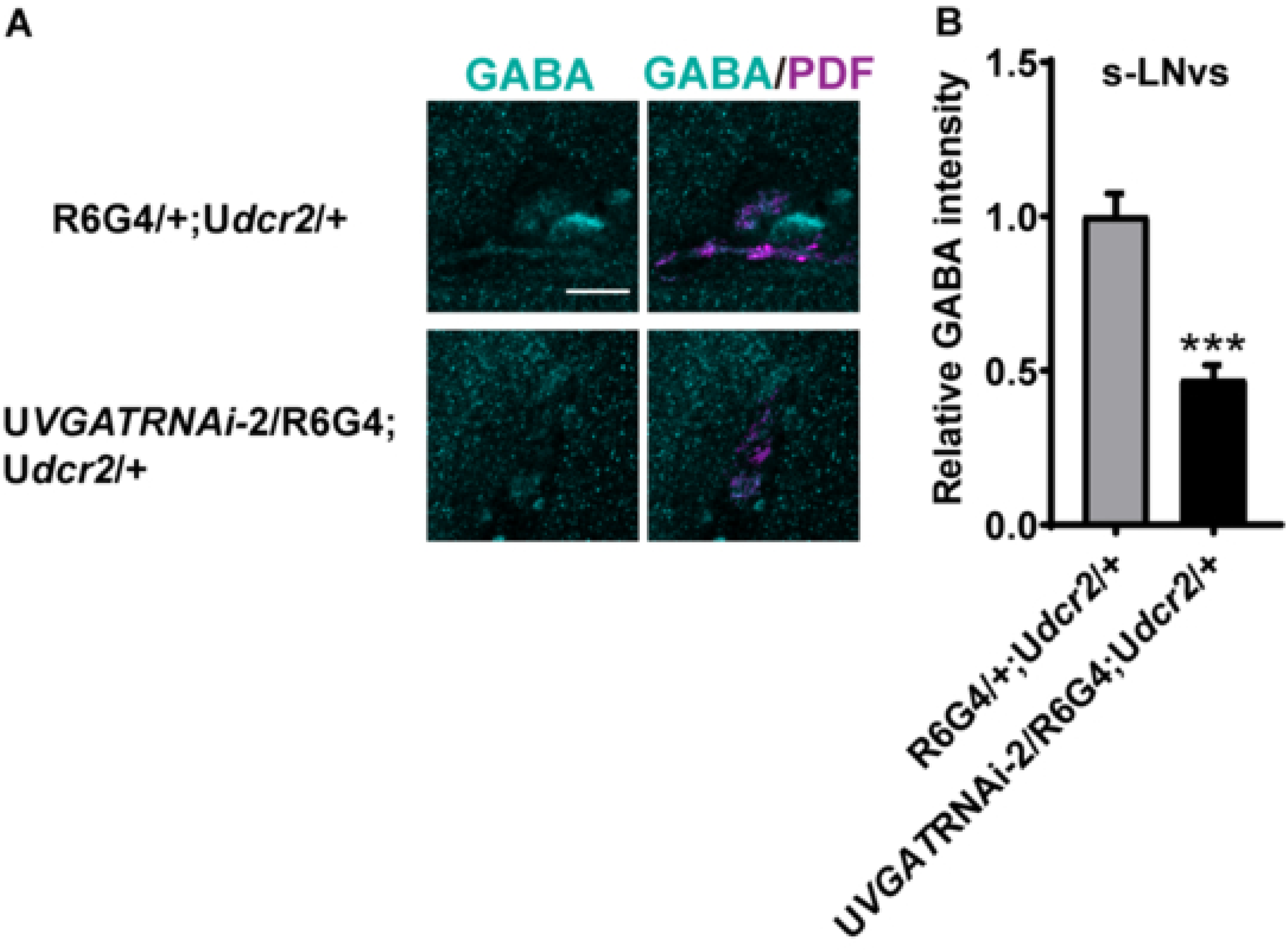
s-LNvs appear to be GABAergic neurons. (A) Brains from male flies with *VGAT* knocked down in the s-LNvs using R6GAL4 and control maintained under 4L20D and immunostained with GABA (cyan) and PDF (magenta) antisera. Representative s-LNvs are displayed. Merged signal is shown as yellow. (B) Bar graphs represent normalized GABA intensity in the s-LNvs with *VGAT* knocked down using R6GAL4 (n = 25, 32 cells). The average value of the control group is set to 1. Two-tailed Student’s *t*-test, \*\*\**P* < 0.001. The scale bar represents 15 µm. Error bars represent SEM. G4, GAL4; U, UAS.

**Figure 5—figure supplement 4.**
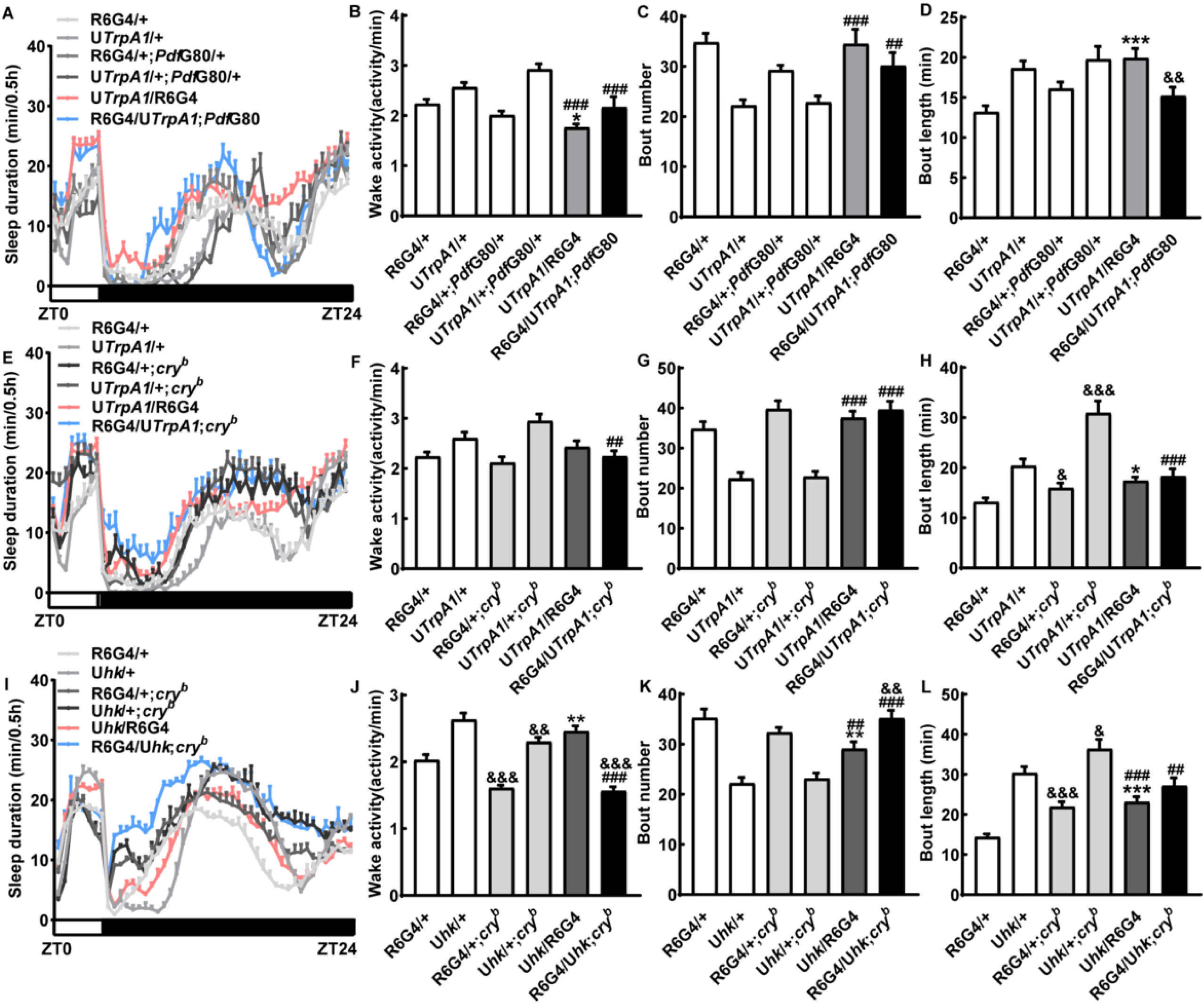
CRY promotes wakefulness by impinging on the excitability of the s-LNvs. (A-D) Sleep profile (A), daily waking activity (B), sleep bout number (C), average sleep bout length (D) of male flies expressing TrpA1 in the s-LNvs using R6GAL4 and relevant controls in Fig 5J, monitored under 4L20D and 29℃ to activate TrpA1. White box indicates light period while black box indicates dark period. (E-H) Sleep profile (E), daily waking activity (F), sleep bout number (G), average sleep bout length (H) of male *cry* mutant flies expressing TrpA1 in the s-LNvs using R6GAL4 and relevant controls in Fig 5K, monitored under 4L20D and 29℃ to activate TrpA1. (I-L) Sleep profile (I), daily waking activity (J), sleep bout number (K), average sleep bout length (L) of male *cry* mutant flies over-expressing HK in the s-LNvs using R6GAL4 and relevant controls in Fig 5L, monitored under 4L20D. For comparison with UAS/GAL4 controls, one-way ANOVA with Bonferroni multiple comparison test was used: compared to GAL4 control, \**P* < 0.05, \*\**P* < 0.01, \*\*\**P* < 0.001; compared to UAS control, ##*P* < 0.01, ###*P* < 0.001. For comparison between mutant vs. control, two-tailed Student’s *t*-test was used: compared to WT background, &*P* < 0.05, &&*P* < 0.01, &&&*P* < 0.001. Error bars represent SEM. G4, GAL4; G80, GAL80; U, UAS.

**Figure 6—figure supplement 1.**
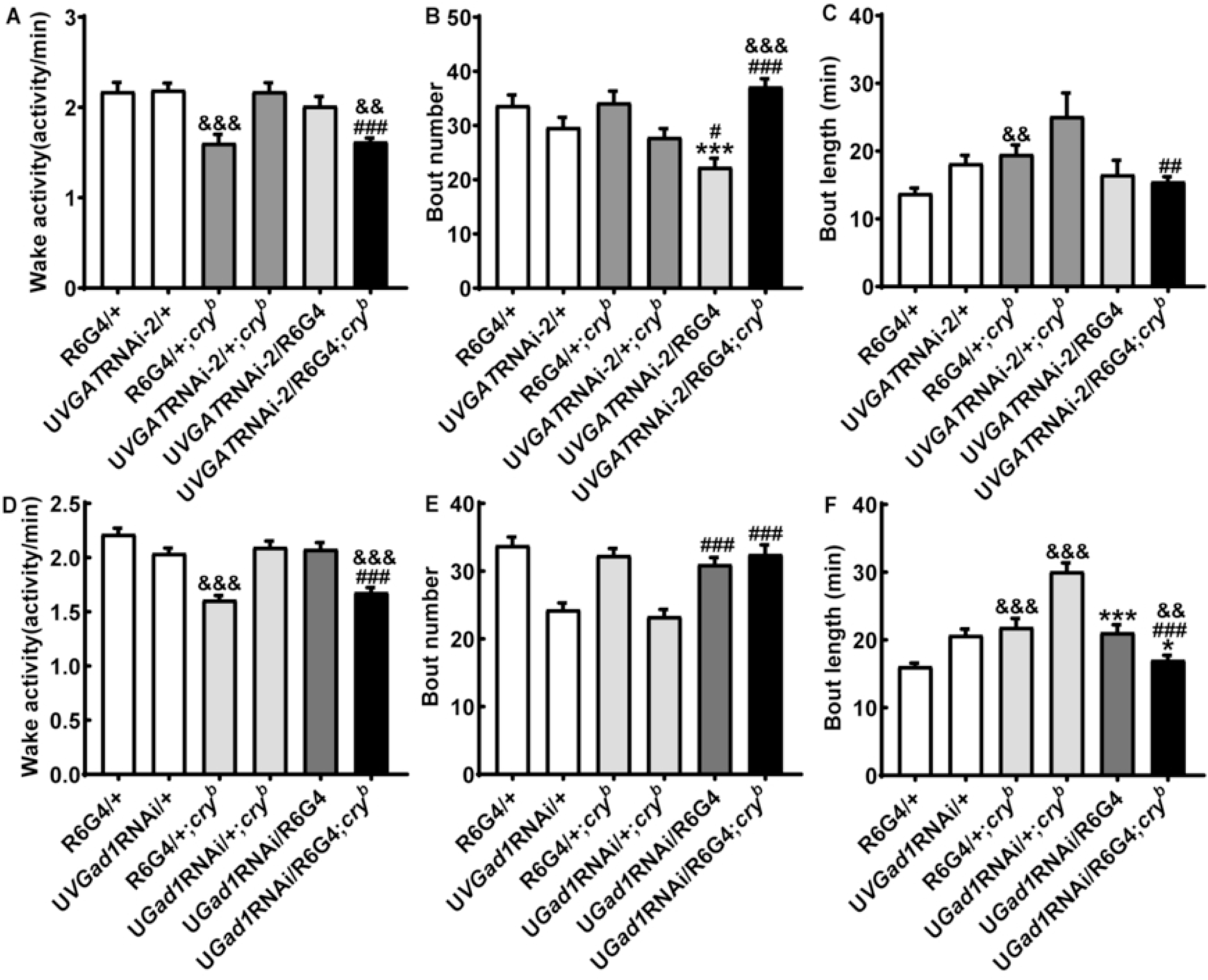
Knocking down *VGAT* or *Gad1* suppress the long-sleep phenotype of *cry* mutation. (A-C) Daily waking activity (A), sleep bout number (B), average sleep bout length (C) of male *cry* mutant flies with *VGAT* knocked down in the s-LNvs using R6GAL4 and relevant controls in Fig 6F, monitored under 4L20D. (D-F) Daily waking activity (D), sleep bout number (E), average sleep bout length (F) of male *cry* mutant flies with *Gad1* knocked down in the s-LNvs using R6GAL4 and relevant controls in Fig 6H, monitored under 4L20D. For comparison between RNAi flies vs. UAS/GAL4 controls, one-way ANOVA with Bonferroni multiple comparison test was used: compared to GAL4 control, \*\*\**P* < 0.001; compared to UAS control, #*P* < 0.05, ##*P* < 0.01, ###*P* < 0.001. For comparison between mutant vs. control, two-tailed Student’s *t*-test was used: compared to WT background, &&*P* < 0.01, &&&*P* < 0.001. Error bars represent SEM. G4, GAL4; U, UAS.

### Source Data

Figure 1—source data 1

Numerical data with associated statistical analyses underlying Figure 1.

Figure 2—source data 1

Numerical data with associated statistical analyses underlying Figure 2.

Figure 3—source data 1

Numerical data with associated statistical analyses underlying Figure 3.

Figure 4—source data 1

Numerical data with associated statistical analyses underlying Figure 4.

Figure 5—source data 1

Numerical data with associated statistical analyses underlying Figure 5.

Figure 6—source data 1

Numerical data with associated statistical analyses underlying Figure 6.

Figure 7—source data 1

Numerical data with associated statistical analyses underlying Figure 7.

